# Physics-informed Deep Learning Approach to Quantification of Human Brain Metabolites from Magnetic Resonance Spectroscopy Data

**DOI:** 10.1101/2022.10.13.512064

**Authors:** Amir M Shamaei, Jana Starcukova, Zenon Starcuk

## Abstract

**Purpose:** While the recommended analysis method for magnetic resonance spectroscopy data is linear combination model (LCM) fitting, the supervised deep learning (DL) approach for quantification of MR spectroscopy (MRS) and MR spectroscopic imaging (MRSI) data recently showed encouraging results; however, supervised learning requires ground truth fitted spectra, which is not practical. Moreover, this work investigates the feasibility and efficiency of the LCM-based self-supervised DL method for the analysis of MRS data.

**Method:** We present a novel DL-based method for the quantification of relative metabolite concentrations, using quantum-mechanics simulated metabolite responses and neural networks. We trained, validated, and evaluated the proposed networks with simulated and publicly accessible in-vivo human brain MRS data and compared the performance with traditional methods. A novel adaptive macromolecule fitting algorithm is included. We investigated the performance of the proposed methods in a Monte Carlo (MC) study.

**Result:** The validation using low-SNR simulated data demonstrated that the proposed methods could perform quantification comparably to other methods. The applicability of the proposed method for the quantification of in-vivo MRS data was demonstrated. Our proposed networks have the potential to reduce computation time significantly.

**Conclusion:** The proposed model-constrained deep neural networks trained in a self-supervised manner can offer fast and efficient quantification of MRS and MRSI data. Our proposed method has the potential to facilitate clinical practice by enabling faster processing of large datasets such as high-resolution MRSI datasets, which may have thousands of spectra.

**Highlights:**

- A novel self-supervised deep learning method for quantifying metabolite concentrations in MR spectroscopy signals.
- Providing a unique opportunity to quantify complex-valued MRS data in the time domain.
- Faster MR spectroscopy quantification with comparable accuracy to traditional methods.
- Investigating the impacts of the dataset size and neural network design on our proposed model

## 1 Introduction

Accurate spectral analysis of in vivo magnetic resonance spectroscopy (MRS) data is complicated by overlapping of resonance lines from different metabolites, which precludes or inhibits accurate identification of low-concentration metabolites, and also by the presence of a broad background of rapidly decaying macromolecule (MM) and lipid signals (Kreis et al., 2021; Robin A. de Graaf, 2019).

Linear combination model (LCM) fitting, peak fitting, and peak integration are the three most often used methods for spectral analysis of an MRS signal (Near et al., 2020a). In LCM fitting, each metabolite in the spectrum is represented by a “basis spectrum,” which is a description of the spectral shape of an individual metabolite (Near et al., 2020a; Poullet et al., 2008; Robin A. de Graaf, 2019; Stagg and Rothman, 2013). Contrary to LCM fitting, peak fitting uses a simple lineshape model function to fit isolated peaks within a spectrum. Peak fitting is highly dependent on the prior knowledge of the parameters of peaks, and imposing a large amount of prior knowledge may be burdensome in crowded spectra like ^1^H-MRS of the brain due to the excessive number of metabolites and peaks per metabolite (Near et al., 2020a; Poullet et al., 2008; Robin A. de Graaf, 2019; Stagg and Rothman, 2013).

LCM fitting is the expert-recommended approach (Near et al., 2020b) owing to its shown efficacy, adaptability, and relative simplicity of usage. The choice of basis spectra over individual peak components (peak fitting method) reduces the number of model functions necessary to adequately describe the spectrum, which leads to fewer fitting parameters. Basis spectra are realistic since they are derived directly from phantom experiments or numerical simulations (Landheer et al., 2019; Near et al., 2020a; Poullet et al., 2008; Robin A. de Graaf, 2019; Stagg and Rothman, 2013; Starcuk and Starcuková, 2017)

A long-standing interest in LCM fitting has resulted in the development of a variety of fitting techniques, including time-domain and frequency-domain algorithms (Daniel G.Q. Chong et al., 2011; Clarke et al., 2021; Oeltzschner et al., 2020; Poullet et al., 2008; Provencher, 2001; Ratiney et al., 2005; Soher et al., 2011; Wilson et al., 2011).

The recent success of deep learning (DL), one of the latest machine learning approaches, in a variety of tasks, including applications with a low signal-to-noise ratio (SNR) (Goodfellow et al., 2016; Lundervold and Lundervold, 2019), suggests that it might also handle the spectral analysis of an MRS signal.

Moreover, supervised DL-based approaches have been used for ghosting artifacts detection and removal (Kreis and Kyathanahally, 2018), spectral reconstruction (Lee et al., 2020), automatic peak picking (Klukowski et al., 2018), magnetic resonance spectroscopic imaging (MRSI) spatial resolution enhancement (Iqbal et al., 2019), localized correlated spectroscopy acceleration (Iqbal et al., 2021), frequency and phase correction of MRS signals (Shamaei et al., 2023; Tapper et al., 2021), and poor-quality spectra identification (Gurbani et al., 2018).

These studies showed results competitive with traditional methods using supervised learning, in which the input and the output were simulated spectra and known values, respectively. The true output values are unknown in in-vivo MRS data. Moreover, a network trained in a supervised manner using simulated data might be prone to overfit training data (Higham and Higham, 2019); thus, any discrepancy between the in-vivo and the simulated training spectra, such as the presence of nuisance peaks, frequency, and phase shifts, and line-broadening, may result in errors in metabolite quantification. Self-supervised learning may eliminate the drawbacks of supervised learning.

A DAE is typically designed to encode the input into a low-dimensional embedding space and then reconstruct the input from the encoding in a self-supervised manner (Goodfellow et al., 2016). A deep autoencoder (DAE) has also been successfully used for metabolite and MM separation in MRS signals, as well as for quantification and noise removal of MRSI signals (Lam et al., 2020; Li et al., 2020).

A common belief among traditional statisticians and DL practitioners is that *’“more data is always better”* (Nakkiran et al., 2019). The effects of model complexity, dataset size, and data augmentation were empirically investigated in computer vision and natural language processing (Belkin et al., 2019; Nakkiran et al., 2019); however, these effects have not been investigated in MRS data quantification using DL.

## 2 Authors’ Contributions

In this paper, we present a novel DL-based LCM algorithm for MRS data quantification, which uses the advantages of LCM and the capabilities of DAE in self-supervised learning. We designed a DAE network that can learn in a self-supervised manner to fit a linear combination of quantum-mechanically simulated basis spectra to the acquired MR spectrum. We utilize the LCM fitting and integrate it into a DAE to estimate relative concentrations of metabolites. We trained, validated, and evaluated our proposed network using simulated MRS data, and in-vivo MRS data obtained from the publicly accessible Big GABA repository (Mikkelsen et al., 2019, 2017). We compared the performance of our proposed model with other LCM packages, namely, QUEST (Ratiney et al., 2005) and FiTAID (Daniel G.Q. Chong et al., 2011), in terms of accuracy and time efficiency.

In addition, we designed an experiment to evaluate the effect of the dataset size and neural network architecture in our proposed model.

## 3 Related Work

Hiltunen et al. (Hiltunen et al., 2002) have demonstrated the feasibility of constructing a quantifying analyzer for long echo time (TE) in vivo proton magnetic resonance spectroscopy (1H NMR) spectra using artificial neural networks with magnitude spectra

Bhat et al. (Bhat et al., 2006) investigated the application of a radial basis function neural network (RBFNN) for the automatic quantification of short echo time, multi-voxel, and phased spectral data.

Hatami et al. (Hatami et al., 2018) and Lee et al. (Lee and Kim, 2019) applied supervised DL approaches to metabolite quantification and presented results comparable to conventional LCM approaches. Chandler et al. (Chandler et al., 2019) also applied a supervised DL approach to study metabolite quantification in edited human brain MRS spectra.

Gurbani et al. (Gurbani et al., 2019) presented a self-supervised DL architecture that integrates a convolutional neural network (CNN) with peak fitting. In their approach, a DAE, introduced by Hinton et al. (Hinton and Salakhutdinov, 2006) as a specific type of neural network, is used as a framework for unsupervised learning.

Shamaei et al. (Shamaei et al., 2021) extracted characteristics from the MRS data using a wavelet scattering transform and predicted metabolite concentrations using a shallow neural network implementation.

Rizzo et al. (Rizzo et al., 2022) explored the effect of DL architectures, spectroscopic input types, and learning designs on the quantification of simulated MR spectroscopy. A list of related work is provided in Table 1.

**Table 1.**
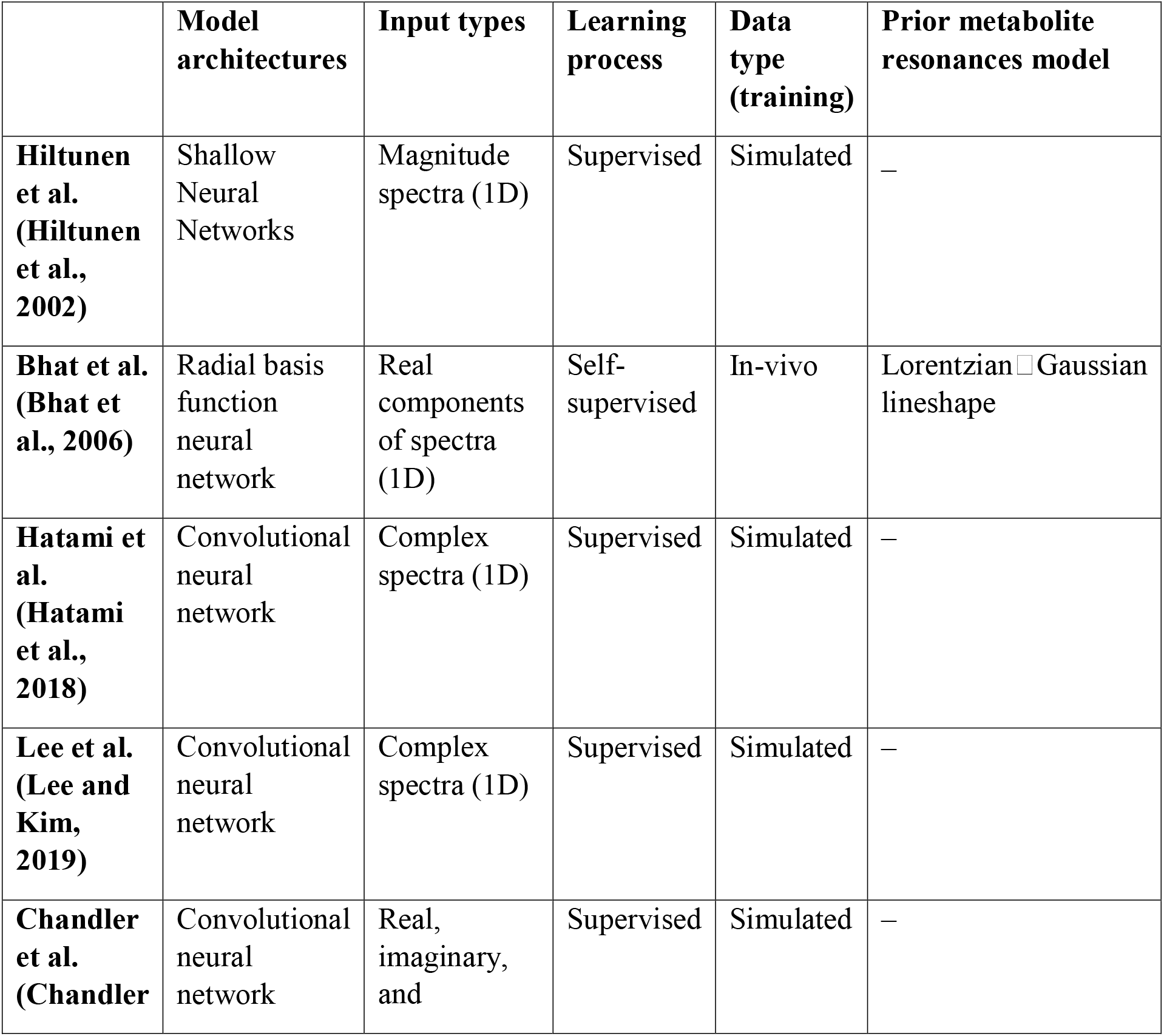
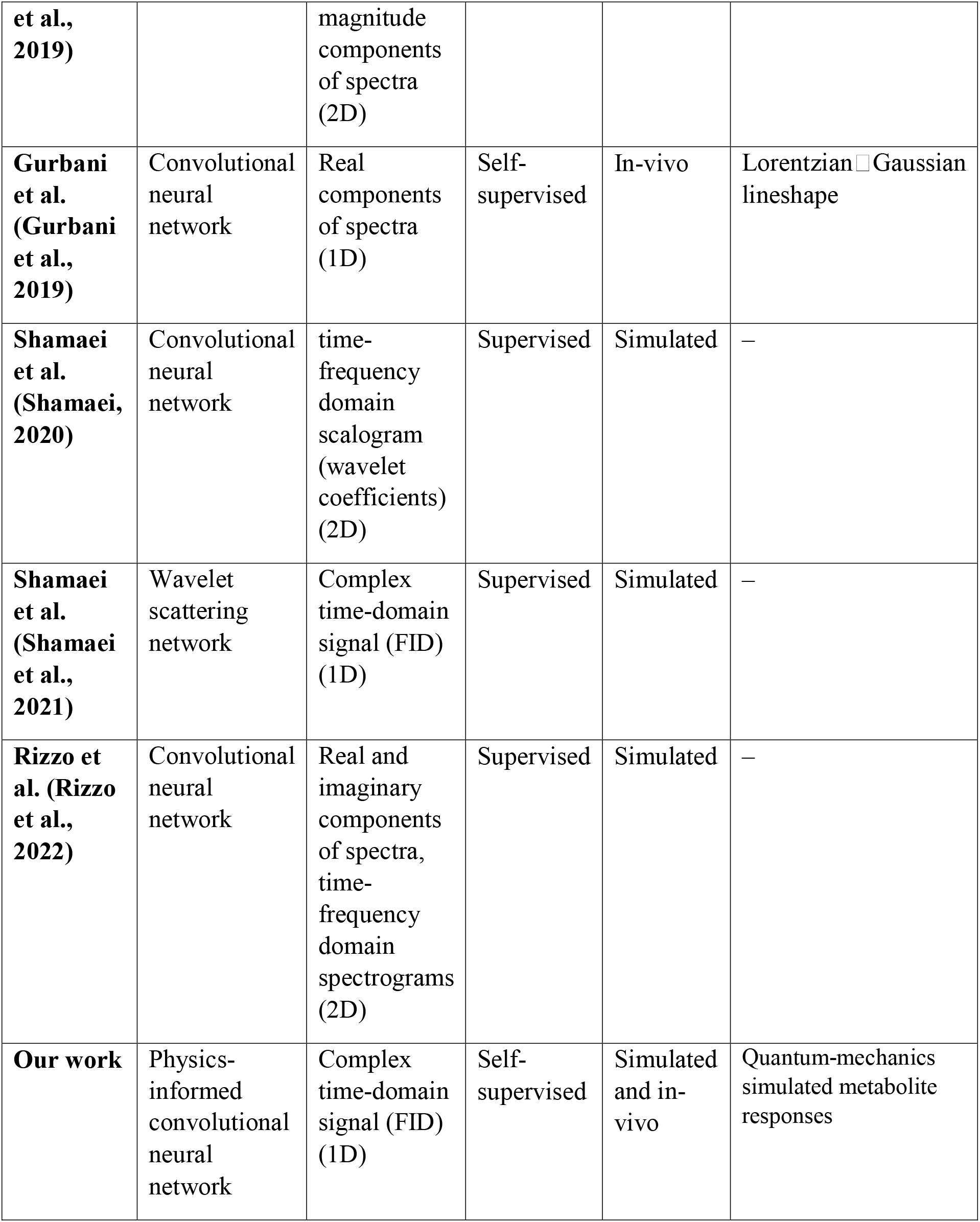
A summary of related work.

## 4 Material and methods

### 4.1 Data Sets

#### 4.1.1 The simulated dataset

One significant challenge in metabolite quantification is that the ground truth values are unknown; thus, it is hard to compare the results of different quantification methods. This problem can be addressed by creating a dataset with known ground truth values using prior physical knowledge and basis spectra.

The MRS signal can be described as a linear combination of amplitude-scaled metabolite signals, the baseline, and noise. The simplified model describing a time-domain MRS signal S(t) as a combination of several metabolite profiles is:

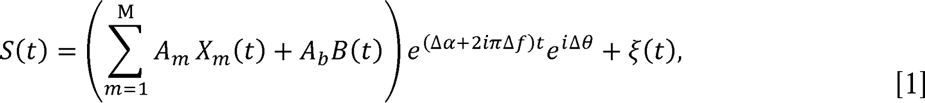

where 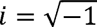 and *A_m_* and *X_m_(t)* are the scaling factor (amplitude) and the model (basis) function (signal) for the *m-th* metabolite, respectively. *Δa*, *Δf*, and *Δθ* are the global damping factor, the global frequency shift, and the global phase shift, respectively. M is the number of metabolites. *A_b_* and £?(t) is the scaling factor and the signal for MMs, and ξ(t) is noise.

Our simulated dataset, containing 96000 signals, was generated using Eq. 1 and the publicly available metabolite basis set from the ISMRM MRS study group’s fitting challenge (Marjańska et al., 2022) (19 metabolites signals (see Table 2) and one MM signal, 3T, PRESS, TE = 30 ms, spectral width = 4000 Hz, 2048 points). The range of parameters *(A_m_, Δa*, *Δf*, and *Δθ)* were determined according to the literature (Govindaraju et al., 2000; Robin A. de Graaf, 2019) (listed in Table 2). The parameters were chosen randomly from the defined ranges with a uniform distribution. The mean SNR of the dataset (largest metabolite peak height in the magnitude spectrum to standard deviation of noise in the frequency domain) was set to 67 ±15 by introducing random complex Gaussian white noise (ξ). Signals in the simulated dataset were shuffled randomly, and 90% of each dataset was allocated to the training subset, 9% to the validation subset, and the rest 1% to the test subset.

**Table 2.** Parameters used in the simulated dataset. *A*_(.)_, Δα, *Δf*, and *Δθ* are the metabolite amplitudes, the global damping factor, the global frequency shift, and the global phase shift, respectively.

| Parameter | Min | Max |
| --- | --- | --- |
| $A_{(\text{Ala})}$ | 0.1 | 1.8 |
| $A_{(\text{Asc})}$ | 0.2 | 1.8 |
| $A_{(\text{Asp})}$ | 1 | 2 |
| $A_{(\text{Cr})}$ | 4.5 | 10.5 |
| $A_{(\text{GABA})}$ | 1 | 2 |
| $A_{(\text{Glc})}$ | 1 | 2 |
| $A_{(\text{Gln})}$ | 3 | 6 |
| $A_{(\text{Glu})}$ | 6 | 12.5 |
| $A_{(\text{GPC})}$ | 0.5 | 1.8 |
| $A_{(\text{GSH})}$ | 1.5 | 3 |
| $A_{(\text{Gly})}$ | 0.2 | 1 |
| $A_{(\text{Ins})}$ | 4 | 9 |
| $A_{(\text{Lac})}$ | 0.2 | 1 |
| $A_{(\text{NAA})}$ | 7.5 | 17 |
| $A_{(\text{NAAG})}$ | 0.5 | 2.5 |
| $A_{(\text{PCho})}$ | 0.2 | 1 |
| $A_{(\text{PCr})}$ | 3 | 5.5 |
| $A_{(\text{PE})}$ | 1 | 2 |
| $A_{(\text{sIns})}$ | 0.2 | 0.5 |
| $A_{(\text{Tau})}$ | 3 | 6 |
| $A_b$ | 14 | 17 |
| $\Delta\alpha$ ($\text{s}^{-1}$) | 0 | 15 |
| $\Delta f$ (Hz) | -3 | 3 |
| $\Delta\theta$ (°) | $-\pi/10$ | $+\pi/10$ |

#### 4.1.2 The in vivo dataset

Data from the public repository Big GABA (Mikkelsen et al., 2019, 2017) were used to demonstrate the applicability of the proposed model for the quantification of in-vivo signals.

We selected 48 single-voxel short-TE PRESS brain datasets (subjects) from 48 healthy volunteers acquired on Siemens scanners from 4 different sites (S1, S5, S6, and S8, 3 Tesla field strength, spectral width = 4000 Hz, 4096 points, TE = 35 ms, TR = 2000 ms; 64 transients; 30 × 30 × 30 mm^3^; medial parietal lobe voxel). Water reference scans without water suppression were obtained with similar parameters, except for 8– 16 averages. It is well recognized that the sufficient size and diversity of data are crucial for the success of the majority of DL models (Goodfellow et al., 2016). However, rich and adequate datasets are uncommon (Lam et al., 2020) in the field of MRS and MRSI. Data augmentation is a possible technique for simulating realistic data by making slight modifications to a small existing dataset.

A two-step data augmentation was applied. In the first step, 16 transients of 64 transients of each subject dataset were randomly selected. Then, the transients were frequency- and phase-aligned using the spectral registration method (Near et al., 2015; Simpson et al., 2017) and were processed into an averaged signal. Eddy-current effects in the averaged signal were corrected using the corresponding water reference signal. Then, the residual water components were removed from the averaged signal with the Hankel Lanczos singular value decomposition (HLSVD) method (Cabanes et al., 2001). The first step was repeated 64 times, producing 64 signals for each subject and thus producing a new extended subject dataset. (Cabanes et al., 2001)

Totally, 3072 (48 × 64) signals, each representing a corrected 16-transient average, were generated and stacked together to create a new in vivo dataset from all subjects.

In the second step of augmentation, first, each complex signal S(t) from the new in vivo dataset generated in step 1 was rescaled as

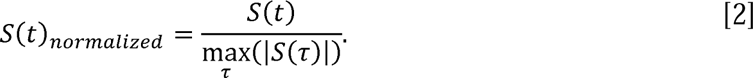

Then five extended subject datasets (320 (5 × 64) signals) were included in a test dataset and excluded from the in vivo dataset. Signals with substantial lipid contamination were removed from the in vivo dataset in the following way: Time-domain signals were Fourier-transformed, then the mean of the magnitude of each signal between 0 and 1.85 ppm was calculated. Finally, based on prior testing on the dataset, signals with a mean value above six were labeled as strongly contaminated signals and discarded from the dataset. Further, we denote this dataset as the filtered dataset (2624 signals).

In the following, ten signals were generated from each signal of the filtered dataset, such that signals were apodized by dampings corresponding to Lorentzian linewidths drawn from a uniform distribution over [0 Hz, 2 Hz]. Then artificial frequency and phase offsets were drawn from a uniform distribution in the ranges of −3 to 3 Hz and −9° to 9°, respectively, and added to apodized signals (ranges were extracted from in vivo brain spectra subjected to spectral fitting). The final training dataset consisted of 28864 (10 × 2624 + 2624) signals. The mean SNR of the training dataset was set to 256 ± 72 by introducing random complex Gaussian noise with a uniform spectral power. Finally, all signals were dedicated to the training set.

To decrease the complexity of the network (number of weights) and the contribution of noise, which typically predominates in the later part of time domain signals (FIDs), the FIDs were shortened to maintain the first 1024 points.

Example spectra from the simulated dataset and the in-vivo dataset are provided in Figure 1.

**Figure 1.**
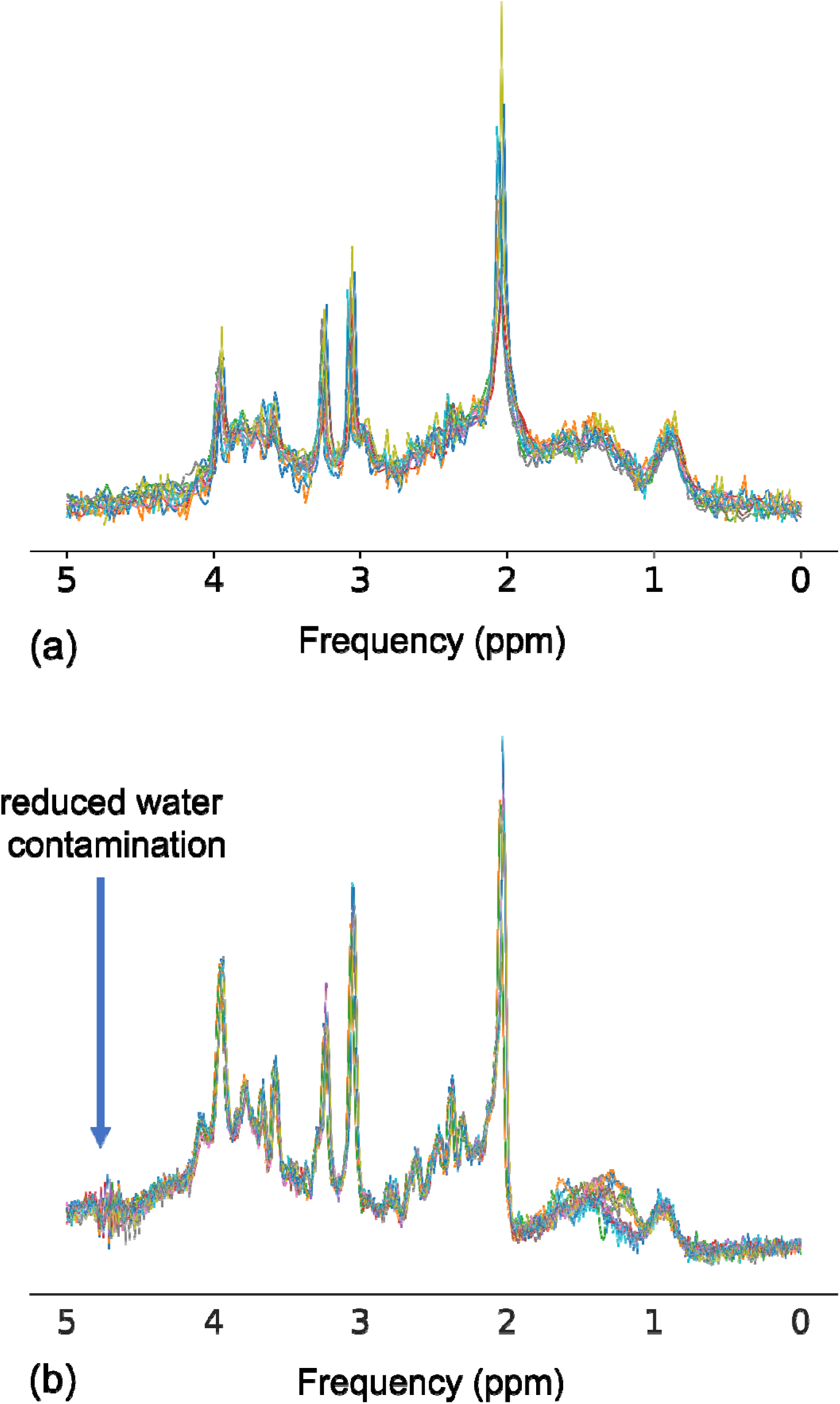
Example spectra from (a) the simulated dataset and (b) the in-vivo dataset.

### 4.2 The deep model

#### 4.2.1 The proposed DAE

DAE is a sort of deep artificial neural network designed to learn an efficient data coding in a self-supervised way (Goodfellow et al., 2016; Hinton and Salakhutdinov, 2006). The basic idea behind DAEs is to utilize the input data as the target that should be reconstructed in the output layer (Goodfellow et al., 2016). A DAE is typically composed of two parts: an encoder and a decoder. The encoder function ***h.*** *= f(x)* maps the n-dimensional input vector (***x*** ɛ *R^n^)* to the n’-dimensional latent vector (***h*** ɛ *R^n^’*), while the decoder function 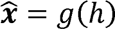 aims to reconstruct the n-dimensional output vector 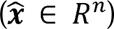 from the latent space representation. Typically, the latent space representation has significantly lower dimension than the input *(n’ « n)*.

Interestingly, the LCM approach toward the quantification of MRS signal can be seen as a DAE, where the few parameters of a model are the interpretable latent space parameters, and the metabolites basis set and prior knowledge are the decoders. For the DAE to be usable for quantifying MRS signals, its latent-space representation must be interpretable as metabolite concentrations. Therefore, it is natural to desire that it includes parameters such as all individual relative metabolite concentrations (amplitudes), damping factors, frequencies, and phases. However, a DAE with the typical architecture (Hinton and Salakhutdinov, 2006) is not useful for quantifying MRS signals.

To this end, a convolutional encoder–model decoder (Gurbani et al., 2019) was employed, incorporating a parametric analysis into a DL model. Our proposed DAE consisted of a conventional encoder and a model decoder. The encoder was composed of sequential layers of convolutional layers (Fukushima, 1980; Schmidhuber, 2015), Gaussian error linear units (GELU) (Hendrycks and Gimpel, 2020), and a fully connected (FC) layer (Goodfellow et al., 2016; Schmidhuber, 2015). Furthermore, a Softplus function (Goodfellow et al., 2016) was applied to estimated amplitudes to ensure that the relative metabolite concentrations (amplitudes) were non-negative. The model decoder was composed of a model function (Eq. 1) that reconstructs the representation of the input signal using the assigned latent space parameters and basis spectra. The proposed DAE architecture is depicted in Figure 2a. The encoder computes four types of parameters: amplitude, frequency, phase, and damping. The input and output of the proposed DAE were complex signals in the time domain.

**Figure 2.**
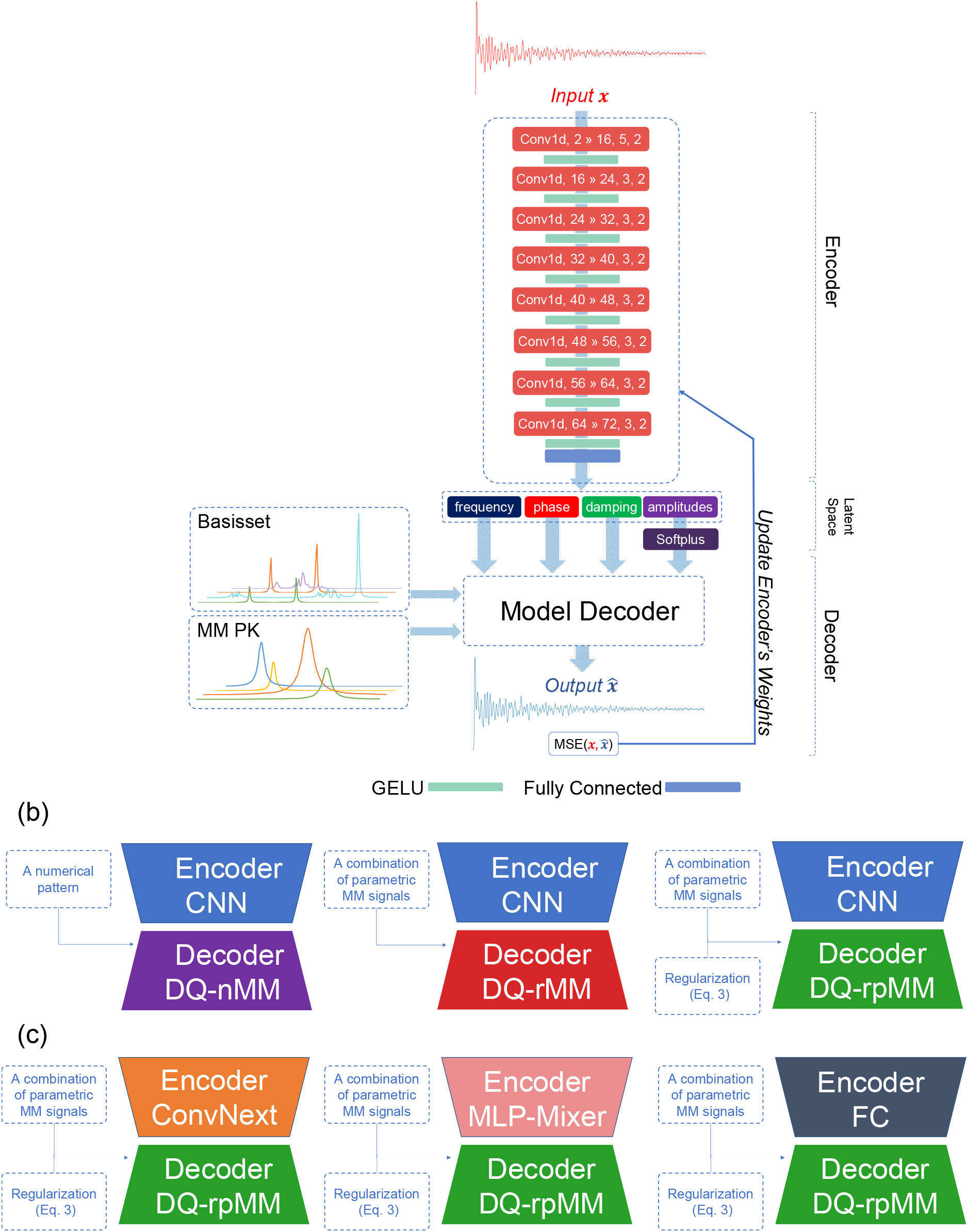
(a) Illustration of the proposed convolutional encoder-model decoder network. The input of the network is a complex signal (x) in the time domain, which is fed to the encoder. The encoder consisted of eight convolutional blocks and an FC layer (see details in Supporting Information Table S2). A convolutional block is composed of a ID convolution (Convld) layer followed by a GELU layer. The model decoder reconstructs the output signal (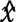) using Eq. 1, basis set, and MM prior knowledge. The DAE was trained to encode the input vector *x* in the time domain into parameters (the amplitudes, damping factor, resonance frequency, and zero-order phase of the basis spectra) that can be used to reconstruct the output vector *it* in the time domain. The proposed network is trained by minimizing the mean square error (MSE) between *x* and 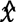. (b) Illustration of three proposed models (DQ-nMM, DQ-pMM, and DQ-rpMM). (c) Illustration of three variants of the DQ-rpMM model with different encoder architectures (ConvNext, MLP-Mixer, and our proposed network with FC layers). CNN, Convolutional Neural Network; Conv1d, (in channels » out channels, kernel size, stride); FC, Fully Connected; GELU, Gaussian Error Linear Unit; MM PK, Macromolecule prior knowledge; DQ-nMM, deep learning-based quantification using a numeric MM; DQ-pMM, deep learning-based quantification using parametric MM; DQ-rpMM, deep quantification using parametric MM and a regularization term.

Training our proposed network is a self-supervised learning task that does not require ground truth values. It minimized the differences between the original input and the consequent reconstruction. In each iteration step of training, the parameters of the encoders were adjusted according to the gradient of the loss function with respect to the given parameters of the model *(A_m_,A_b_*, Δα, Δƒ, and Δθ). The mathematical representation of training can be written as follows:

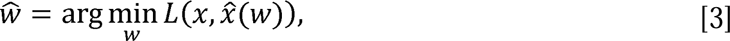

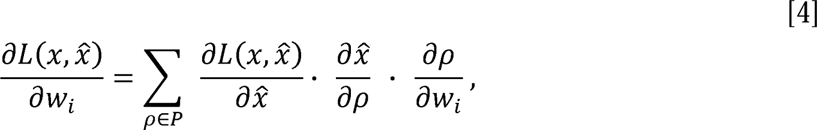

where **x** and 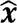 are the n-dimensional input vector 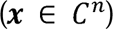 and the n-dimensional output vector 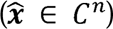, respectively. Eq. 3 is the objective function of training, *L* is the loss function, *w_t_* is an element of the parameters set (weights and biases) of the encoder, and 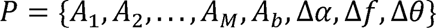 Gradients were computed with the help of Pytorch using its automatic gradient computation (Paszke et al., 2019), which is a reverse automatic differentiation system.

#### 4.2.2 Modeling of macromolecules

Short-TE MRS signals contain a contribution of MMs that overlap with metabolites (Cudalbu et al., 2021). Thus, MM modeling or MM removal from the signal should be included in the spectra analysis.

The presence of MMs may be accounted for in a variety of ways (Near et al., 2020b). MMs can be included as a numerical pattern (found, for instance, from metabolite-nulling inversion recovery measurements) (D G Q Chong et al., 2011). Another approach is to include a set of parametric patterns in the spectral fitting algorithm (Near et al., 2020a; Robin A. de Graaf, 2019).

We implemented both approaches to MM handling in the proposed DAE, which resulted in two models. The first model (referred to as deep learning-based quantification using a numeric MM pattern [DQ-nMM]) includes a MM signal as a numerical pattern (D G Q Chong et al., 2011) in Eq. 1. The second model (referred to as deep learning-based quantification using parametric MM components [DQ-pMM]) includes a MM signal as a combination of parametric MM components in Eq. 1 (the detailed model is provided in Supporting Information Text S1).

However, the increased number of fitted parameters without constraints increases the risk of over-parametrization [42] in the DQ-pMM model. Another approach to handling MM is to remove the initial part of the FID (Poullet et al., 2008), which eliminates MM signals thanks to their decay being faster than that of metabolites. Nevertheless, the disadvantages of this approach are neglecting important information in the early part of the time domain signal and difficulty in determining the number of removed initial points (Poullet et al., 2008).

To overcome these disadvantages and handle the over-parametrization problem in the DQ-pMM model, we proposed a novel approach for MM modeling, which resulted in the third model (referred to as deep learning-based quantification using parametric MM components and a regularization term [DQ-rpMM]).

DQ-rpMM includes a combination of parametric MM components in Eq. 1 and adds a regularization term (Eq. 5) to the loss function.

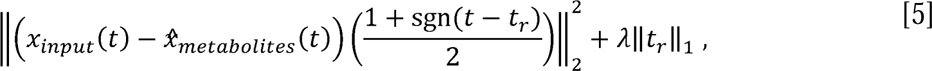

where sgn is a sign function, its shift *t_r_* is a parameter determined by the network, *x_input_* and 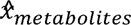 are the input signal and the reconstructed signal from the metabolite components, respectively. Parameter *λ* tunes the significance of the regularization term. Brackets 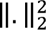 and 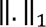 represent squared L2-norm and L1-norm, respectively.

The first term promotes consistency between 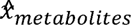 and *x_input_.* Intuitively, the cutoff time *t_r_* should increase to a point where the contribution of MM is low, and the contribution of metabolites is reasonably high. Excessive cutoff of useful signal (that would lead to fitting the non-informative end tail of 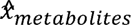 to the noisy part of *x_input_)* should be prevented by the contradictory force of the second term. The architectures of the proposed models are depicted in Figure 2b.

### 4.3 Implementation Details and Training

All steps were run on a computer with a dual EPYC 7742 (2×64 cores) processor and one graphics processing unit (NVIDIA A100 40 GB). The DAE was implemented in Python programming language (Paszke et al., 2019) with the help of the Pytorch lightning interface (Lightning, 2020).

The initial architecture of the network and the training parameters were optimized using the Bayesian Optimization HyperBand algorithm (Falkner et al., 2018) with the help of the Tune framework (Liaw et al., 2018). The optimization details are given in Supporting Information Text S2. All training was performed using the mean-squared error loss (MSE) and the Adam optimizer (Kingma and Ba, 2015) with a batch size of 128, a learning rate of 0.001, and 200 epochs. An early-stopping strategy (Paszke et al., 2019) was performed by monitoring the MSE of the validation subset at the end of every epoch and stopping the training when no improvement was observed in 50 epochs. As reported (Loshchilov and Hutter, 2016), reducing the learning rate is beneficial when learning becomes stagnant. The learning was reduced twice by a factor of 10 during training. Our source codes and data (in NIfTI-MRS format (Clarke et al., 2022)) are available at https://github.com/isi-nmr/Deep-MRS-Quantification.

All proposed models (DQ-nMM, DQ-pMM, and DQ-rpMM) were tested on the simulated dataset. Moreover, The DQ-rpMM model was tested on the in vivo dataset to show the applicability of our method to in-vivo data in which a numeric pattern of MMs is not available. For the simulated dataset, the basis set and the numeric pattern of MMs from the ISMRM MRS study group’s fitting challenge (Marjańska et al., 2022) were utilized in the model decoder. The provided numeric pattern of MMs was parametrized using AMARES (Vanhamme et al., 1997) algorithm and utilized in the models with parametrized MMs, i.e., DQ-pMM and DQ-rpMM (the details of the parametrization are provided in Supporting Information Table S1). For the in vivo dataset (Big GABA), the publicly available metabolite basis set (Zöllner et al., 2021) consisted of 19 metabolites was used in the model decoder: alanine (Ala), aspartate (Asp), creatine (Cr), negative creatine methylene (-CrCH2), γ-aminobutyric acid (GABA), glycerophosphocholine (GPC), glutathione (GSH), glutamine (Gln), glutamate (Glu), water (H2O), myo-inositol (mI), lactate(Lac), NAA, N-acetylaspartylglutamate (NAAG), phosphocholine (PCho), PCr, phosphoethanolamine, scyllo-inositol, and taurine. MM components (13 gaussian lineshapes, previously reported in (Birch et al., 2017)) were used to generate the lipid and MM basis signals.

### 4.4 Accuracy analysis (for the simulated data set)

For the simulated data set, in which the true amplitudes were known, we compared the results, i.e., the estimated amplitudes, obtained with our approach with those obtained by traditional LCM methods using mean absolute percentage error (MAPE) and the coefficient of determination (R^2^) metrics. It has been reported that R^2^ is more informative than MAPE (Chicco et al., 2021). In fact, MAPE does not reveal much information about the effectiveness of regression with regard to the distribution of the ground truth values.

### 4.5 Monte Carlo analysis

Monte Carlo (MC) studies were carried out to investigate the bias and the standard deviation of the proposed DQ-nMM estimates. A signal without noise was generated using Eq. 1 with known global frequency (2 Hz), phase (45°), damping (5 Hz), and known amplitudes of metabolites and MMs. Second, 256 realizations of a normally distributed random complex noise were added to the signal such that SNR was in the range of ∼50.

Finally, the amplitudes of metabolite components were estimated for the 256 signals using the network trained with the simulated dataset, and the estimation errors (MAPE) were measured.

### 4.6 Data size effects

In this experiment, five variants of the simulated dataset with different sizes (1000, 12000, 24000, 96000, and 384000 signals) were generated using the same procedure mentioned for generating the simulated dataset. 90% of each dataset was allocated to the training subset and 10% to the validation subset. Then, the first proposed model (DQ-nMM) was trained using each training dataset. The performance of each dataset in terms of R^2^ of each metabolite and the reconstruction loss (MSE) was monitored using its validation subset during training.

### 4.7 The regularization term effects

The significance of the regularization term (i.e., Λ) in Eq. 5 determines from which time point, 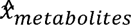 and *x_input_* must be consistent. An experiment was designed to address the effect of *λ* on the performance of the proposed model. DQ-rpMM methods were trained seven times with varying *λ* ({0,1,2,3,4,6,81). All runs used the simulated dataset with a size of 24000 (90% of each dataset was allocated to the training subset and 10% to the validation subset) and were trained for 65000 iterations. All other training parameters were identical to those indicated in the Implementation Details section.

### 4.8 The architecture effects

To test the applicability of successful machine-vision architectures, we used two cutting-edge architectures, ConvNext (Liu et al., 2022) and MLP-Mixer (Tolstikhin et al., 2021), as the encoder of the DQ-rpMM model, which resulted in two variants of the DQ-rpMM model.

To test the effect of convolutional layers on the performance of our proposed models, we substituted FC layers for the convolutional layers in the DQ-rpMM model, which resulted in the third variant of the DQ-rpMM model.

We modified ConvNext (Liu et al., 2022) and MLP-Mixer (Tolstikhin et al., 2021) architectures to process 1D signals. The architectures of the proposed variants are depicted in Figure 2c. The details of ConvNext, MLP-Mixer, and our proposed network with FC layers architectures are provided in Supporting Information Table S3, S4, and S5.

These three variants of the DQ-rpMM model were trained for 100 epochs using the simulated dataset and the parameters mentioned in section 2.4 for DQ-rpMM, except that the initial learning was decreased to 1 ×10**’**^4^

Table 3 gives a list of the experiments that were carried out in this study.

**Table 3.** A list of experiments.

| **Experiment** | **Dataset** | **Models** |
| --- | --- | --- |
| **Monte Carlo analysis** | simulated | DQ-nMM |
| **data size effects** | simulated | DQ-nMM |
| **Regularization term effects** | simulated | DQ-rpMM |
| **Architecture effects** | simulated | DQ-rpMM(CNN),<br>DQ-rpMM(FC),<br>DQ-rpMM(MLP-Mixer),<br>DQ-rpMM(ConvNext), |

## 5 Results and Discussion

All proposed models were validated using 128 signals selected randomly from the test signals of the simulated dataset, and the DQ-rpMM model was evaluated using the test subset of the in vivo datasets (320 signals).

Based on the results and interpretation of the fitting challenge conducted by the MRS study group of ISMRM (Marjańska et al., 2022), in this work, two LCM fitting methods were used, namely, QUEST (Ratiney et al., 2005; Stefan et al., 2009) and FiTAID (Daniel G.Q. Chong et al., 2011), in order to compare the results obtained for the simulated data and determine whether our proposed models are generally comparable to existing fitting procedures.

The purpose of utilizing QUEST was for its optimal performance in the challenge for the artifact-free healthy-brain-like dataset (Marjańska et al., 2022). FiTAID performed close to the top in the challenge (Marjańska et al., 2022) and also offers fitting in both time and frequency domains, allowing for a wider comparison.

The simulated test signals were processed using the QUEST method (the QUASAR plugin in the jMRUI software package (Stefan et al., 2009)) in two distinct ways. The first procedure (labeled as QUEST) included the numeric MM signal in the basis set. In the second procedure (labeled as QUEST-Subtract), the MM signal was not included in the basis set but estimated with the QUEST subtract technique (Ratiney et al., 2005) (the first 100 points were cut off).

The simulated test signals were processed using the FiTAID software in the time (labeled as FiTAID [time]) and frequency (labeled as FiTAID [freq.]) domains over specified ranges (TD = [1..1024]) points, FD = [0,5] ppm). Eq.1 was utilized as a model in the QUEST and the FiTAID software for quantification. Table 4 contains a list of used methods.

**Table 4.** A list of all utilized models in this study.

| Model | Dataset |  | MM model | Fitting domain |
| --- | --- | --- | --- | --- |
|  | simulated | in vivo |  |  |
| **DQ-nMM (ours)** <sup>□</sup> | ✓ | — | numerical pattern | Time |
| **DQ-pMM (ours)** <sup>□</sup> | ✓ | — | parametric MM components | Time |
| **DQ-rpMM (ours)** <sup>□</sup> | ✓ | ✓ | parametric MM components and regularization term |  |
| **FiTAID [time]** <sup>†</sup> | ✓ | — | numerical pattern | Time |
| **FiTAID [freq.]**<sup>†</sup> | ✓ | — | numerical pattern | Frequency |
| **QUEST**<sup>†</sup> | ✓ | — | numerical pattern | Time |
| **QUEST-Subtract**<sup>†</sup> | ✓ | — | subtract<br>technique<br>(Ratiney et al.,<br>2005) | Time |
| □ The source code is available at https://github.com/isi-nmr/Deep-MRS-Quantification<br>† QUEST and FiTAID software are available in the jMRUI software package (version 7) at http://www.jmrui.eu/ |  |  |  |  |

Table 5 and Figure 3 illustrate a comparison of our DL-based proposed models and traditional methods using the test signals of the simulated dataset. On average, DQ-nMM produced better performance in terms of R^2^ and MAPE than QUEST and FiTAID (time and freq.). QUEST-Subtract showed the worst performance in terms of R^2^ and MAPE. DQ-rpMM presented a slightly better performance compared with DQ-pMM in all terms. The DQ-pMM method with parametrized MM showed poor performance.

**Figure 3.**
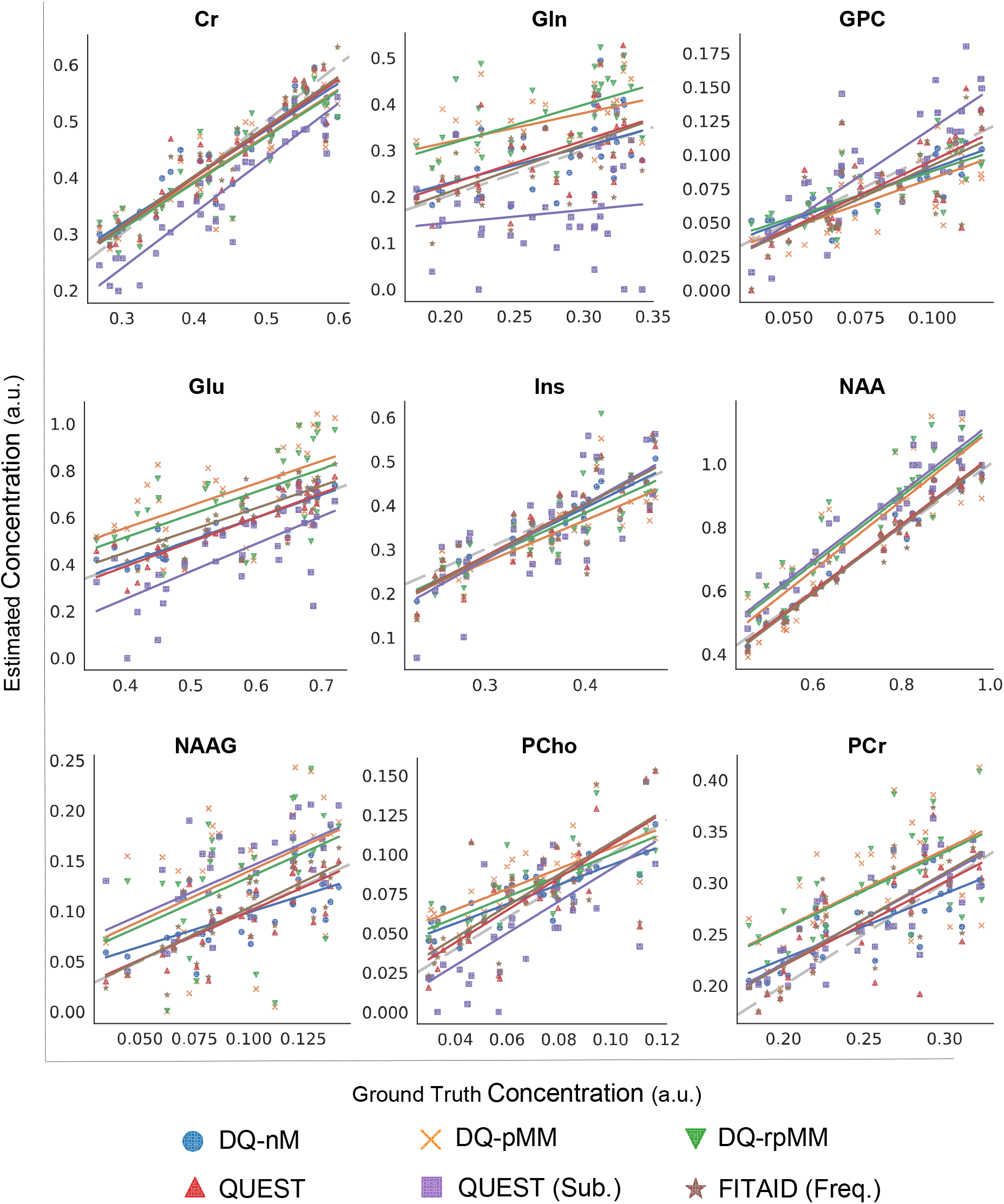
Testing the results of the simulated dataset. Plots of estimated concentrations against the ground truth concentrations in arbitrary units (a.u.). Algorithms are color-coded. DQ-nMM, deep learning-based quantification using a numeric MM; DQ-pMM, deep learning-based quantification using parametric MM; DQ-rpMM, deep quantification using parametric MM and a regularization term; Quest (Sub.), Subtract-QUEST; FiTAID (freq.), FiTAID with frequency domain fit.

**Table 5.**
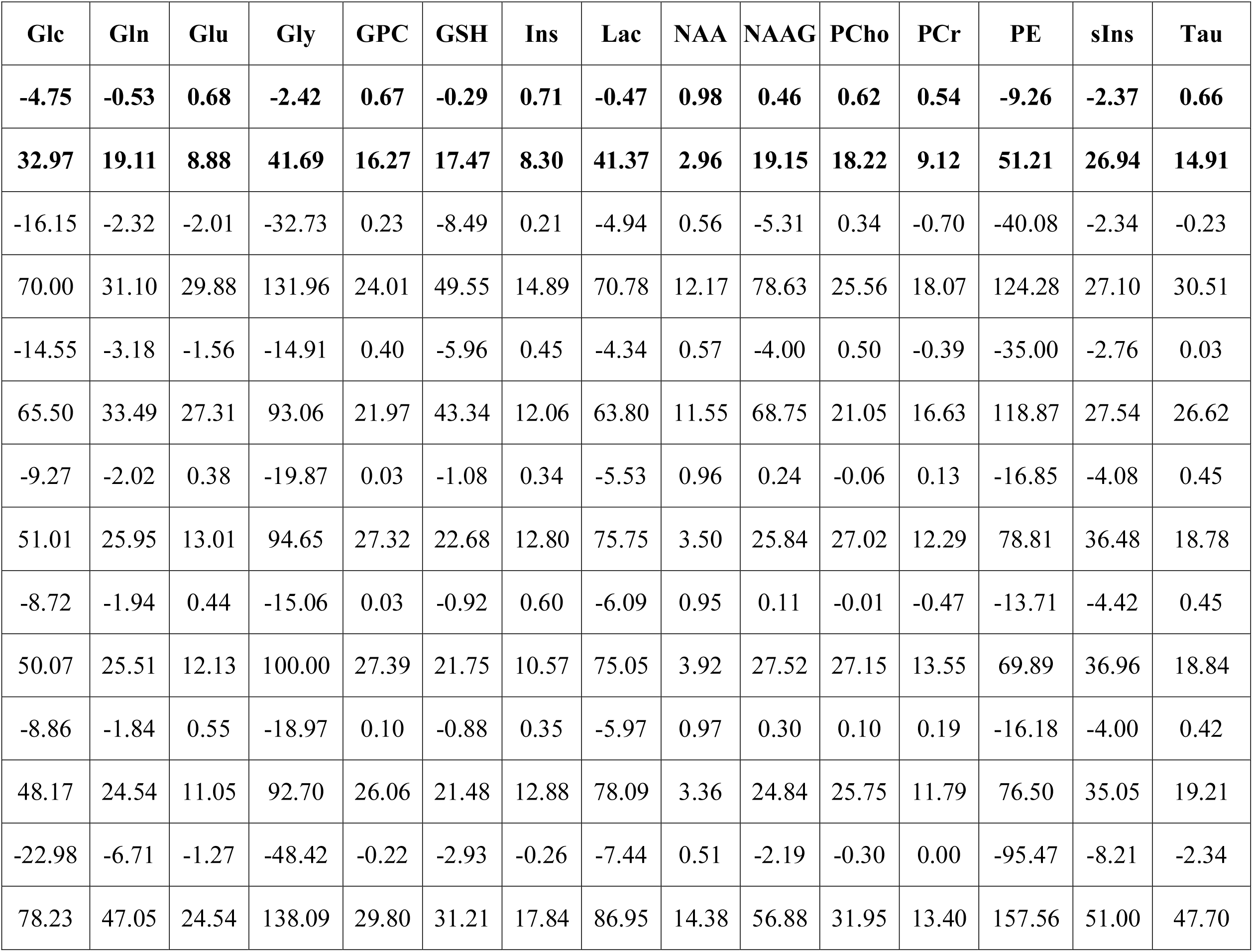
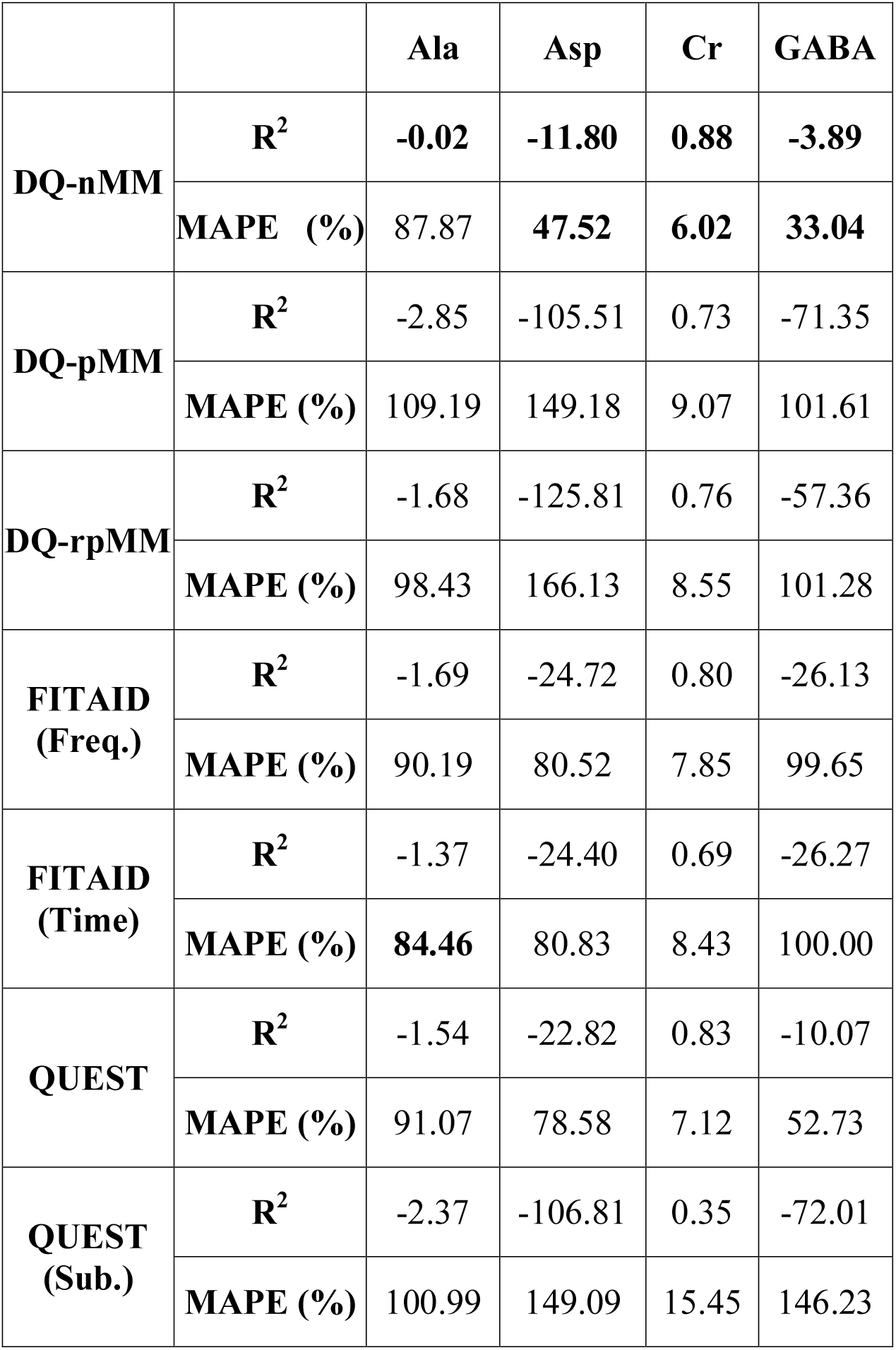
Testing results of the simulated dataset using DQ-nMM, DQ-pMM, DQ-rpMM, QUEST, and FiTAID. R^2^ and MAPE are calculated between the true and estimated values. All metabolites and seven methods are reported in rows and columns, respectively. MAPE, mean absolute percentage error; R^2^, coefficient of determination; MM, macromolecule; DQ-nMM, deep quantification using numeric MM; DQ-pMM, deep quantification using parametric MM; DQ-rpMM, deep quantification using parametric MM and a regularization term; QUSET (Sub.), Subtract-QUEST; FiTAID (Time), FiTAID with the time-domain fit; FiTAID (freq.), FiTAID with the frequency-domain fit.

DQ-nMM outperformed other methods in the quantification of most metabolite concentrations. DL-based methods showed an excellent performance in the estimation of Cr concentration. However, DQ-pMM and DQ-rpMM struggled to estimate NAA concentration. In general, all methods reported relatively high MAPEs in the estimation of GPC, PCho, Tau, and NAAG concentrations. All methods showed a poor performance in estimating low-concentration metabolites, for example, Ala, Asp, GABA, Gly, and PE. The effect of damping, frequency, phase, and SNR of spectra on quantification accuracy is provided in Supporting Information Figure S1.

The training time for DQ-nMM, DQ-pMM, and DQ-rpMM were 40.7, 73.9, and 89.6 minutes, respectively. The processing time per signal for DQ-nMM, DQ-pMM, and DQ-rpMM were 0.031, 0.507, and 0.517 milliseconds, respectively. The processing time per signal for FiTAID (time), FiTAID (freq.), QUEST, and QUEST-Subtract were 2500, 2500, 560, and 1518 milliseconds, respectively.

Figure 3 illustrates scatterplots between estimated and ground truth values for eight metabolites (Cr, Gln, GPC, Glu, Ins, NAA, NAAG, PCho, PCr, and Tau). The values of R^2^, MSE, and *t_r_* during training are reported in Supporting Information Figure S3. DQ-nMM revealed a very good performance in the estimation of Cr (R^2^=0.88), Glu (R^2^=0.68), GPC (R^2^=0.67), Ins (R^2^=0.71), NAA

(R^2^=0.98), PCho (R^2^=0.62), PCr (R^2^=0.54), and Tau (R^2^=0.66) concentration and a good performance in estimation of NAAG (R =0.46). Interestingly, DQ-nMM outperformed QUEST in the estimation of GPC (R^2^=0.10), Ins (R^2^ =0.35), PCho (R^2^ =0.10), PCr (R^2^ =0.19), and Tau (R^2^ =0.42).

Figure 4 illustrates example spectra from the test signals of the simulated dataset quantified by DQ-nMM, DQ-pMM, and DQ-rpMM, along with the ground truth spectra. On visual inspection, DQ-pMM and DQ-rpMM showed higher residual compared with DQ-nMM.

**Figure 4.**
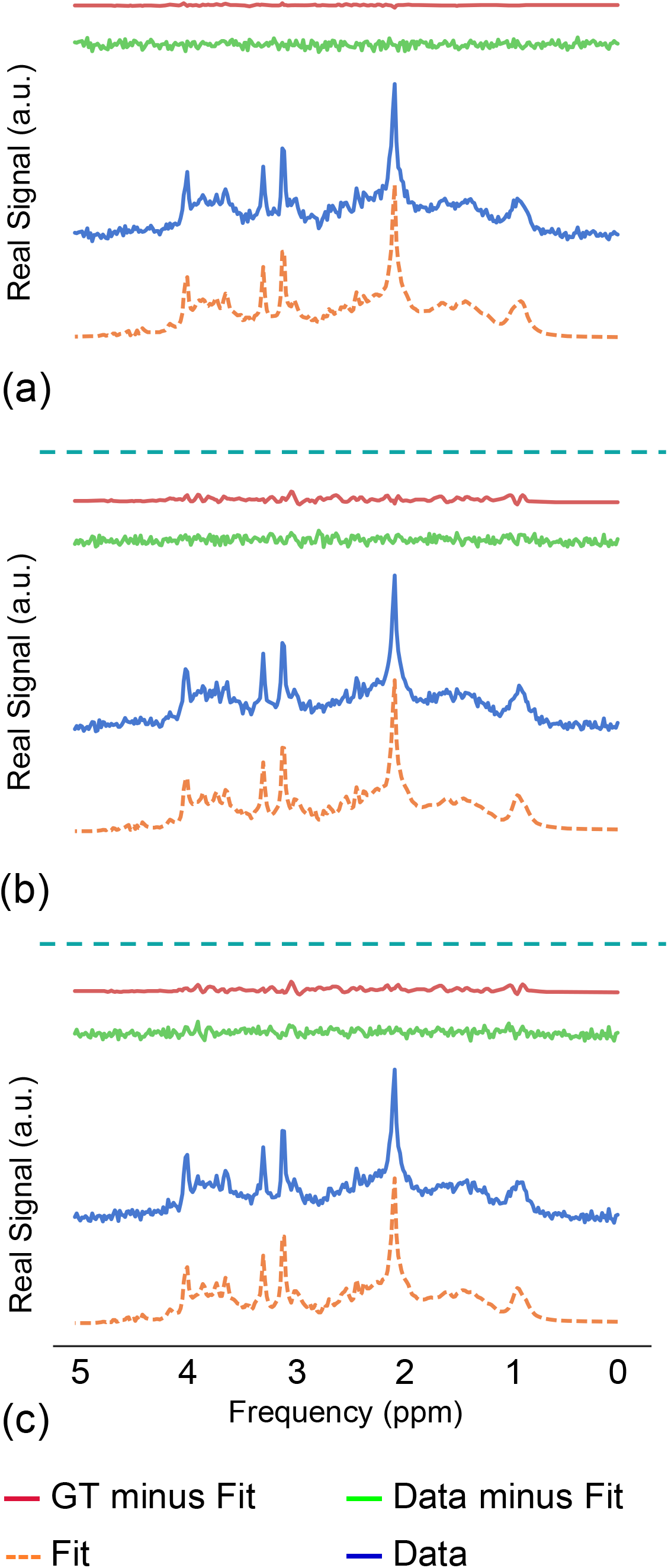
Example spectra from the test subset of the simulated dataset quantified by (a) DQ-nMM, (b) DQ-pMM, and (c) DQ-rpMM. Spectra are color-coded. The contribution of each basis spectrum to the total spectrum was illustrated in Supporting Information Figure S6. GT, ground truth; DQ-nMM, deep learning-based quantification using a numeric MM; DQ-pMM, deep learning-based quantification using parametric MM; DQ-rpMM, deep quantification using parametric MM and a regularization term.

Figure 5 shows the results of the MC analysis for the DQ-nMM method. Figure 5a shows the distribution of estimated concentrations as histogram plots. In most metabolites, the histogram peak is close to the true value, except that the concentration of NAAG was slightly underestimated. Figure 5b shows a scatter visualization of a joint distribution of estimated frequencies and dampings of spectra, illustrating the bias and variance of the DQ-nMM method for individual metabolites in the test subset of MC analysis. Results revealed robust performance in detecting dampings and frequencies of spectra. Figure 5c shows a heatmap of a correlation matrix representing the correlation between estimated concentrations in the test subset of MC analysis. The heat map highlighted a high correlation between PCr and Cr, and GPC and PCho.

**Figure 5.**
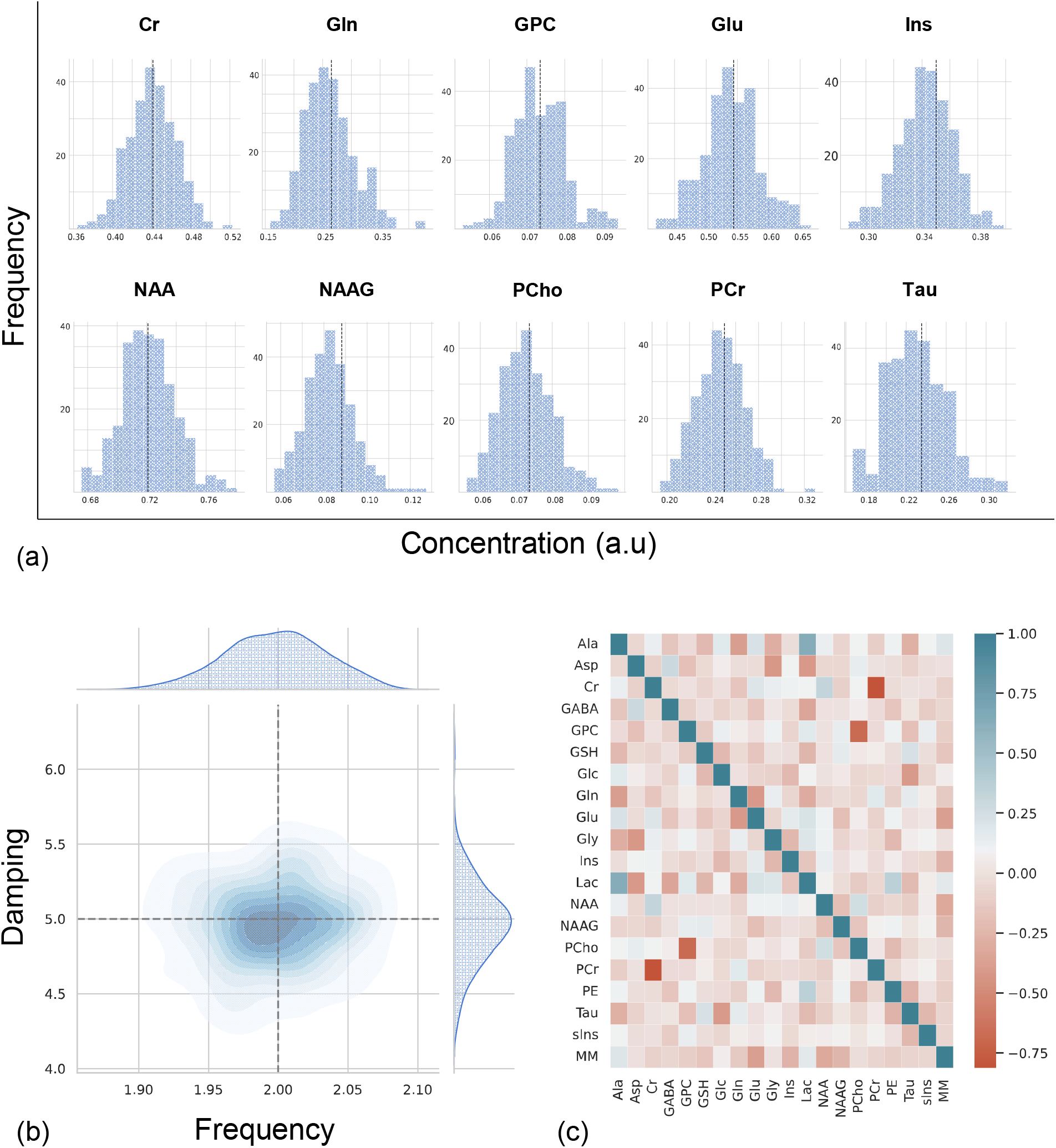
The results of the MC analysis. Spectra are quantified by the DQ-nMM method, (a) Histograms of the estimated concentrations. The vertical lines are true values, (b) A scatter visualization of a joint distribution of estimated frequencies (Hz) and dampings (Hz) of spectra. The dashed vertical and horizontal lines are the true frequency and damping, respectively. (c) A graphical representation of the correlation matrix representing the correlations between estimated concentrations. A strip plot of estimated concentration is provided in Supporting Information Figure S2. DQ-nMM, deep learning-based quantification using a numeric MM; MC, Monte Carlo.

Figure 6a shows the effect of dataset size during the training of the DQ-nMM method using the validation subset. The mean R^2^ value is calculated across selected metabolites (Cr, GPC, Ins, NAA, PCho, Tau). Figure 6b highlights the effect of *λ* on the performance of the DQ-rpMM method. The mean R^2^ value is calculated across the selected metabolites (Cr, GPC, Ins, NAA, PCho, Tau). Moreover, the trend of *t_r_* in the DQ-rpMM method with seven different *λ* during training is shown.

**Figure 6.**
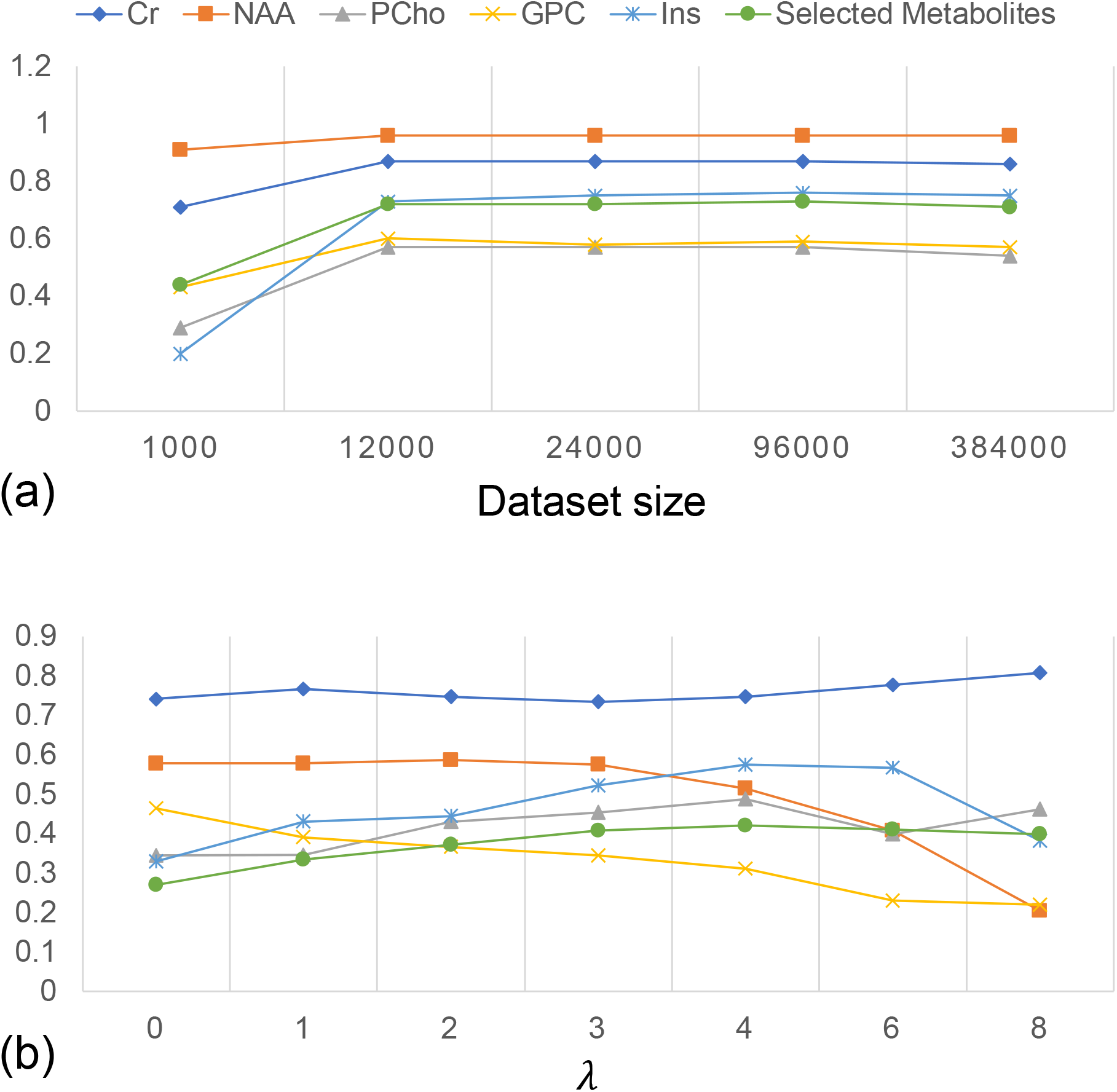
(a) The mean R^2^ value calculated across the selected metabolites (NAA, Cr, PCho, GPC, and Ins), and the R^2^ value for each of the selected metabolites obtained (a) from the DQ-nMM model trained with datasets with sizes of 1000, 12000, 24000, and 96000, and 384000, (b) from DQ-rpMM trained with different *λ.* The figure shows that the choice of *λ* can affect the performance of quantification of a specific metabolite. For instance, lambda with a high value resulted in a good performance in the quantification of PCho, and Cr, and, conversely, a poor performance in the quantification of NAA and GPC. DQ-nMM, deep learning-based quantification using a numeric MM; DQ-rpMM, deep learning-based quantification using parametric MM and a regularization term; R^2^, coefficient of determination; *λ*, the significance of the regularization term.

The influence of the encoder architecture on the performance of the proposed model (DQ-rpMM) is illustrated in Figure 7. The performance of the proposed networks with convolutional layers was superior to those with fully connected layers.

**Figure 7.**
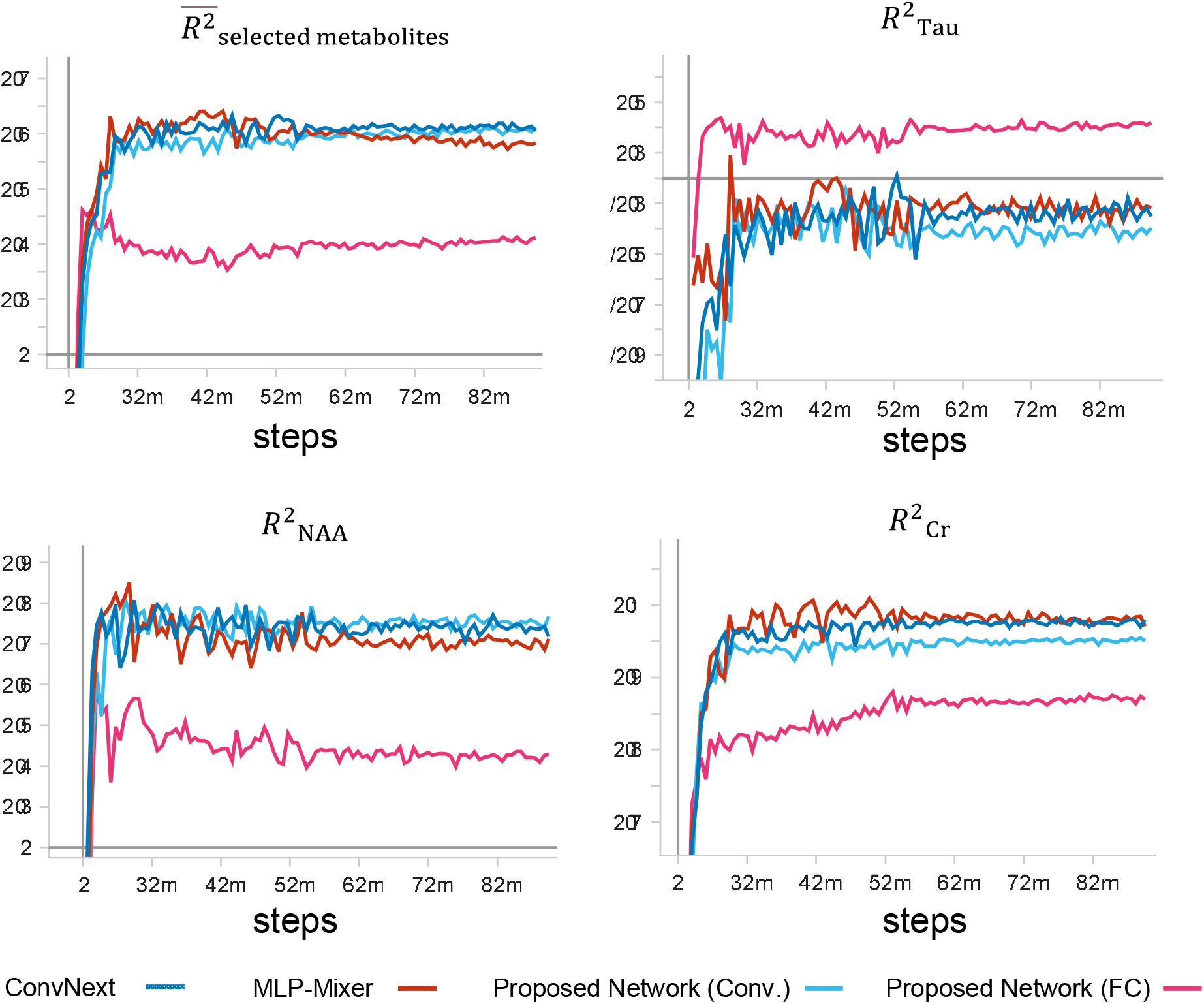
Online monitoring of metrics during training of DQ-rpMM with ConvNext (dark blue), MLP-mixer (red), our proposed convolutional (Conv.) network (light blue), and our proposed fully connected (FC) network (pink). The mean R^2^ value calculated across the selected metabolites (Cr, GPC, Ins, NAA, PCho, Tau) and the R^2^ values for NAA, Cr, and Tau against training steps are plotted. DQ-rpMM, deep learning-based quantification using parametric MM components with a regularization term; R^2^, coefficient of determination.

Figure 8 illustrates four example spectra from the test subset of the in vivo dataset quantified by DQ-rpMM. The reconstruction capability of our proposed model is clearly shown. The mean, standard deviation (SD), and coefficient of variation (CV) of each metabolite-to-creatine ratio for five subjects are summarized in Table 6 for in-vivo data. The CV was lowest for Glx (0.9%) and highest for tCho (10.6%) in subject 3. The means of the tNAA-to-Cr ratio were relatively consistent among subjects. Moreover, the SDs of the tNAA-to-Cr ratio were found to be minimal.

**Figure 8.**
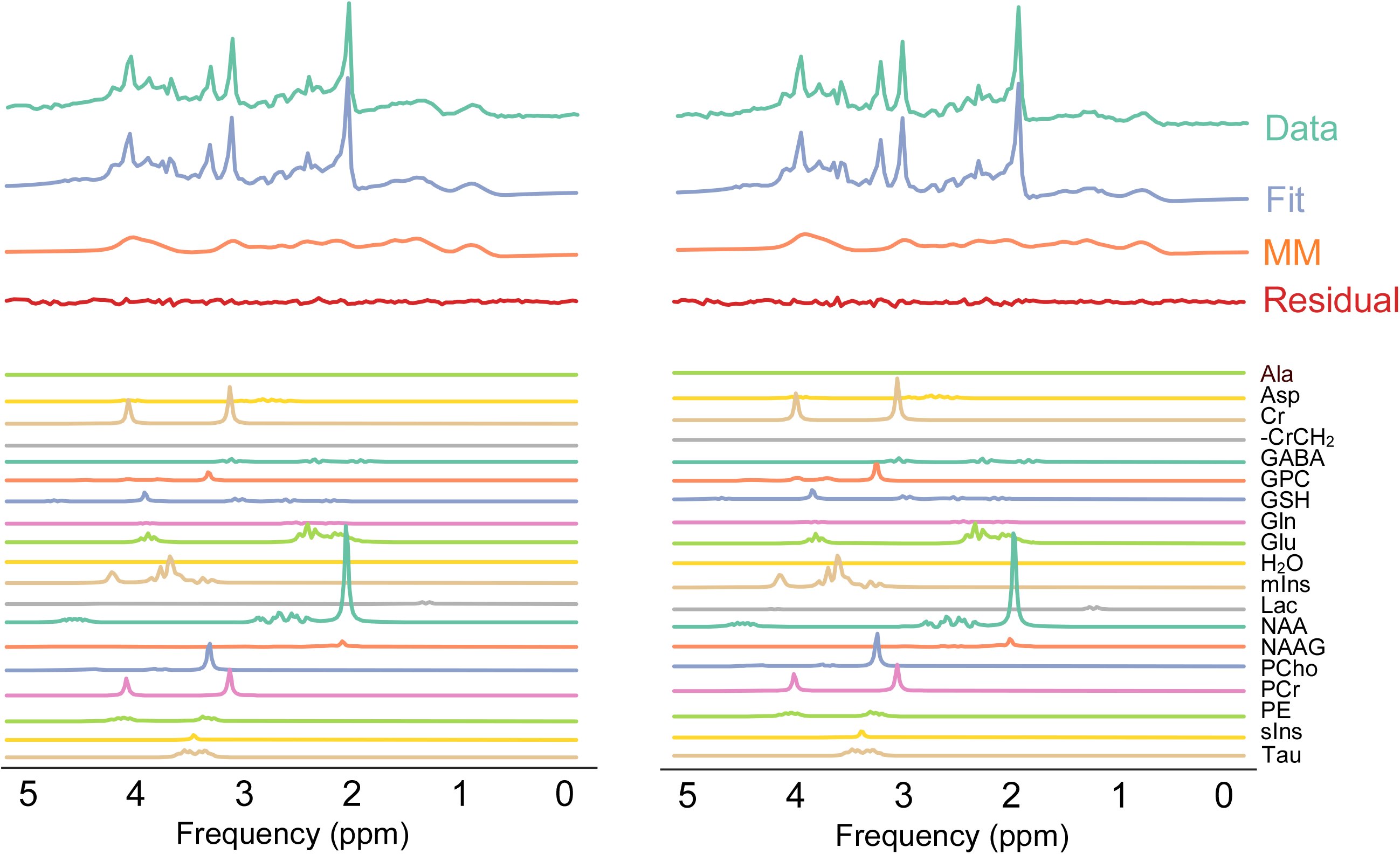
Two example spectra from the test subset of the in vivo dataset quantified by DQ-rpMM. Spectra are color-coded. A strip plot of the estimated relative concentration within subjects and two more example spectra are provided in Supporting Information Figures S4 and S7. GT, ground truth; DQ-rpMM, deep learning-based quantification using parametric MM and a regularization term.

**Table 6.**
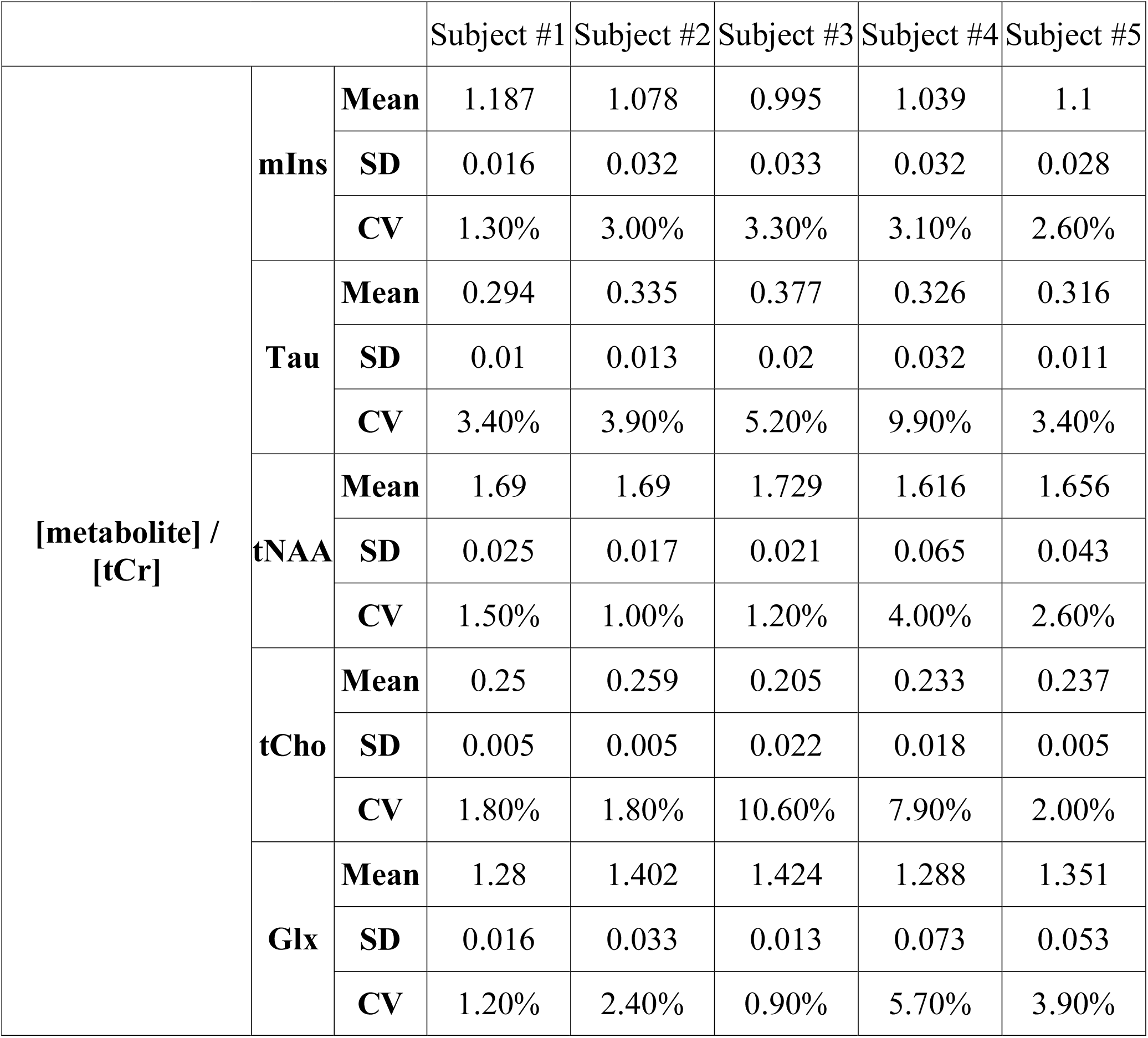
Testing results of the in-vivo dataset using DQ-rpMM. Five metabolites are reported for the five subjects. Each subject has 64 augmented spectra. The mean, standard deviation (SD), and coefficient of variation (CV [SD/mean]) of each metabolite-to-creatine ratio are listed by subjects. DQ-rpMM, deep quantification using parametric MM and a regularization term.

Given the recent agreement on using LCM fitting (Near et al., 2020a) and several pioneering research highlighting the possible use of DL in MRS (Gurbani et al., 2019, 2018; Kreis and Kyathanahally, 2018; Lee and Kim, 2019), this work demonstrates that an LCM fitting-based self-supervised DL approach can be used to quantify both simulated and in-vivo MRS data.

No labels or ground truth were required during training since the proposed method maximized the use of prior knowledge in a deep neural network to limit a model solution. The network can identify the contribution of basis metabolite spectra and MM to crowded MRS spectra such as those in Figure 4 and Figure 8. The network can reconstruct the input signal by a linear combination of basis spectra, therefore highlighting the contribution of each basis spectrum to the total spectrum (Figure 4 and Figure 8). Moreover, the difference between the data and fit can be calculated and served as an indicator of whether the fitting operation was successful or not (Stagg and Rothman, 2013).

The network was validated using simulated data in which in-silico ground truth values were available. A comparison between this novel DL-based LCM and traditional LCM fitting algorithms, namely QUEST, QUEST-Subtract, and FiTAID, was performed (Figure 3, Table 5). The results show that the DL-based algorithm can achieve comparable performance with a significantly shorter amount of time for quantification of an MRS signal (0.031 ms) compared with traditional methods (560 ms).

For metabolites such as NAA, Ins, Glu, Tau, and Cr, which have relatively high concentration levels, all methods provided with a numeric MM pattern (DQ-nMM, QUEST, FiTAID) showed good performance (Figures 3 and Table 5). Noteworthy, DQ-nMM showed remarkable results for metabolites such as PCho (R^2^ = 0.62, MAPE = 18.2 %) and PCr (R^2^ = 0.54, MAPE = 9.12 %) which overlap with GPC (R^2^ = 0.67, MAPE = 16.27 %) and Cr (R^2^ = 0.88, MAPE = 6.02 %), respectively, while traditional method, such as QUEST struggled with quantification of GPC (R^2^ = 0.1, MAPE = 26.06 %), PCr (R^2^ = 0.19, MAPE = 11.79 %) and PCho (R^2^ = 0.10, MAPE = 25.75 %). Moreover, DQ-nMM showed a good performance in the decomposition of NAA and NAAG, which is notoriously challenging owing to the low concentration of NAAG and spectral overlap (Edden et al., 2007).

DQ-rpMM showed better performance than DQ-pMM and QUEST-Subtract. The use of the regularization terms in DQ-rpMM almost doubled performance compared to DQ-pMM in the quantification of metabolites such as GPC and Ins and PCho in terms of R^2^. The reported MAPEs also support our observation (Table 5). Our proposed model DQ-nMM also performed well in an MC analysis where the performance of estimations of metabolites level was found to be very good, and no significant bias was observed (Figure 6).

The main drawback of supervised deep learning quantification of MRS data is (i) the lack of model interpretability and (ii) the need for ground truth data. Introducing prior knowledge and a model function to a deep neural network can provide a self-supervised interpretable deep learning approach to MRS data quantification. Gurbani et al. previously presented a model-informed self-supervised neural network in which the information of peak components was required as prior knowledge; however, their method depends heavily on prior information of the excessive number of metabolites and peaks per metabolite which is time-consuming in particular for spectra acquired in higher magnetic fields. Our approach incorporates realistic quantum-mechanics simulated metabolite responses, eliminating the need for the information of peak components, e.g., the location of NAA resonance, or any other constraints.

The computational bottleneck in processing volumetric MRSI is accurate quantification, partly because the current LCM approaches rely on iterative algorithms. Since our method showed comparable performance compared with QUEST and FiTAID, once trained, our proposed method can simply be implemented on scanner computers, allowing for fast (10000 spectra in 80 seconds using a standard computer) and accurate quantification. As reported in (Simicic et al., 2021), our findings confirm that when a numerical MM pattern is available, methods that are equipped with this type of pattern, such as DQ-nMM, should be utilized due to their good performance.

In general, one of the main weaknesses of DL models is that they cannot perform well on new data acquired using different acquisition parameters. In the case of the same pulse sequence with either longer or shorter TE, it is necessary for our method to be retrained using a new basis set; however, transfer learning methods can mitigate this issue.

Along with demonstrating the performance of the proposed approach on simulated data, the method DQ-rpMM was used to conduct quantification in publicly accessible in-vivo MRS data (Big GABA). Moreover, we demonstrated the generalizability of the proposed model to unseen in-vivo data (Figure 8 and Table 6).

Our results (Table 6) highlighted that CVs for tNAA, mIns, and Glx were consistent among subjects. For tCho and Tau, CVs were less consistent. In general, our proposed model showed stability in the quantification of spectra within subjects.

We observed that the performance of our method could also be affected by the parameters of the network and prior knowledge (Figures 6 and 7). Our results indicate that the accuracy of our proposed method trained on a dataset with 1000 samples is suboptimal, and a training dataset with a minimum size of 12000 samples is necessary for optimal quantification. As reported by Nakkiran et al. (Nakkiran et al., 2019), our results verify that increasing the number of training samples reduces test performance. It is interesting since big datasets are rare in MRS applications.

In the field of machine learning, it is typical to estimate the computation cost using FLOPS (floating-point operations per second) (Liu et al., 2022). The computational cost of our CNN was 2.66 M FLOPS. The computational costs (notated with *O) of* the iterative Levenberg-Marquardt method based on the QR factorization as the linear solver is (Lin et al., 2016): *O(n ×m) + k(O(m^3^) + O(n ×m) + O*(m^3^)) where *n* is the number of data points and *m* is the number of model parameters, and *k* is the number of the Levenberg-Marquardt damping parameters that are being used (Lin et al., 2016). By setting *k* to 1 for the sake of simplicity, *m* to 24, and *n* to 2048, the total cost of Levenberg-Marquardt at every iteration is 1.4 M FLOPS. For example, FiTAID (Daniel G.Q. Chong et al., 2011) requires 100 iterations by default, and the computational cost of a fitting procedure using traditional LCM is above 100 MFLOPS.

We investigated the influence of deep neural network design by comparing our proposed network to existing architectures for machine vision tasks. As depicted in Figure 7, substituting convolutional layers with FC layers resulted in a substantial drop in performance and an increase in computational cost (4.52 MFLOPs).-The results (Figure 7) indicate that MLP-Mixer with extremely deep designs may perform worse. Furthermore, ConvNext demonstrated the same performance as our proposed convolutional network. However, ConvNext and MLP-Mixer demanded greater computational power (FLOPs for ConvNext and MLP-Mixer were 710 MFLOPs and 2.4 GFLOPs, respectively). Our results suggest that (i) networks with simple and shallow design can perform optimal quantification of MRS data and (ii) using convolutional layers are necessary.

Because of using a model-decoder as the decoder, our proposed method can calculate the Cramér-Rao lower bounds to estimate uncertainty. However, additional work is required to extend the capability of the proposed method for quantifying MRS data with simultaneous uncertainty estimation. Additionally, it is important to evaluate how the distribution of parameters in the training set affects the performance of the network.

In this work, we utilized a simplified model for describing a time-domain MRS signal (Eq.1); however, we can expand the capabilities of our proposed DAE to handle more complex models. While the proposed DL-based method for quantification of MR spectroscopy data shows promising results, there are some limitations to consider:

First, the proposed method was validated using simulated and publicly accessible in-vivo human brain MRS data, which may not fully represent the variability and complexity of clinical data. Therefore, further validation with a larger dataset that includes more diverse patient populations is needed to evaluate the generalizability of the method.

Second, the proposed method relies on quantum-mechanics simulated metabolite responses, which may not fully capture the variability of in-vivo metabolite responses. This may result in some inaccuracies in the quantification of metabolite concentrations.

Third, the proposed method is constrained by the LCM model, which may limit its ability to detect and quantify metabolites that are not included in the model. Therefore, the proposed method may not be suitable for the analysis of MRS data that contains metabolites that are not well characterized by the LCM model.

Lastly, the proposed method requires a large amount of training data to optimize the neural network, which may limit its applicability in certain settings where obtaining large datasets is challenging.

## 6 Conclusions

In this study, we presented a self-supervised deep learning technique for accurate quantification of in-vivo and simulated MRS data using linear combination modeling. Our proposed approach eliminates the need for labeled data during training and highlights the contribution of each basis spectrum in the overall spectrum. We compared our method with conventional approaches and observed that it quantified MRS signals with comparable performance but in a significantly shorter amount of time. Moreover, we demonstrated that more complex and deeper architectures did not improve the performance of our shallower architecture.

We proposed a novel approach for MM modeling which increased the performance of our method. Our results indicated that a training dataset with a minimum size of 12000 is required for precise quantification, and larger datasets do not improve quantification accuracy.

In conclusion, our study presents a self-supervised DL-based approach for accurate MRS data quantification, offering a faster alternative to conventional techniques. Our findings could be of significant value in accelerating the quantification of large MRS datasets. Future studies should investigate the clinical relevance of our proposed method.

## 7 List of all Supplementary Information Captions

Supporting Information Figure S1. Scatter plots of the MAPEs in the estimated Cr and NAA concentration versus damping (Hz), frequency (Hz), phase (°), and SNR of spectra.

Supporting Information Figure S2. The results of MC analysis.

Supporting Information Figure S3. Online monitoring of metrics during training.

Supporting Information Figure S4. Strip plot of estimated relative concentration within subjects.

Supporting Information Table S1. The details of the parametrization of the MM signal.

Supporting Information Table S2. Summary of the proposed network (Convolutional).

Supporting Information Table S3. Summary of the proposed network (MLP-Mixer).

Supporting Information Table S4. Summary of the proposed network (ConvNext).

Supporting Information Table S5. Summary of the proposed network (Fully connected).

Supporting Information Figure S6. Example spectra from the test subset of the simulated dataset quantified by (a) DQ-nMM, (b) DQ-pMM, and (c) DQ-rpMM highlighting the contribution of metabolite. Spectra are color-coded.

Supporting Information Figure S7. Four example spectra (a,b,c, and d) from the test subset of the in vivo dataset quantified by DQ-rpMM.

Supporting Information Text S1: The detailed model of macromolecules.

Supporting Information Text S2: Bayesian hyper-parameterization.

## 8 DATA AVAILABILITY STATEMENT

The source code is freely available at [https://github.com/isi-nmr/Deep-MRS-Quantification]. For questions, please contact the authors. The data that support the findings of this study are openly available in Big GABA at https://www.nitrc.org/projects/biggaba/.

## 10 ACKNOWLEDGEMENTS

This work is part of the project that has received funding from the European Union’s Horizon 2020 research and innovation program under the Marie Sklodowska-Curie grant agreement No 813120 (INSPiRE-MED) and was also supported by institutional support RVO:68081731 - Czech Academy of Sciences, Institute of Scientific Instruments.

This work was supported by the staff and equipment of the ISI-MR facility of the Czech-Biolmaging infrastructure, supported by grants CZ. 02.1.01/0.0/0.0/18_046/0016045, LM2018129 and LM2023050 of the MEYS CR.

The authors thank Radim Kořínek, Ph.D. (Czech Academy of Sciences, Institute of Scientific Instruments, Czech Republic), Reza Goodarzi, Rudy Rizzo, and Mohammad Jafarian for their valuable technical support.

