## SUPPLEMENTAL MATERIAL for "Physics-informed Deep Learning Approach to Quantification of Human Brain Metabolites from Magnetic Resonance Spectroscopy Data" (from another version of this preprint: version 2, 2022-10-29)

### Supporting Information

Supporting Information Figure S1. Scatter plots of the MAPEs in the estimated Cr and NAA concentration versus damping (Hz), frequency (Hz), phase (˚), and SNR of spectra.

Supporting Information Figure S2. The results of MC analysis.

Supporting Information Figure S3. Online monitoring of metrics during training.

Supporting Information Figure S4. Strip plot of estimated relative concentration within subjects.

Supporting Information Table S1. The details of the parametrization of the MM signal.

Supporting Information Table S2. Summary of the proposed network (Convolutional).

Supporting Information Table S3. Summary of the proposed network (MLP-Mixer).

Supporting Information Table S4. Summary of the proposed network (ConvNext).

Supporting Information Table S5. Summary of the proposed network (Fully connected).

Supporting Information Text S1: The detailed model of macromolecules.

Supporting Information Text S2: Bayesian hyper-parameterization.

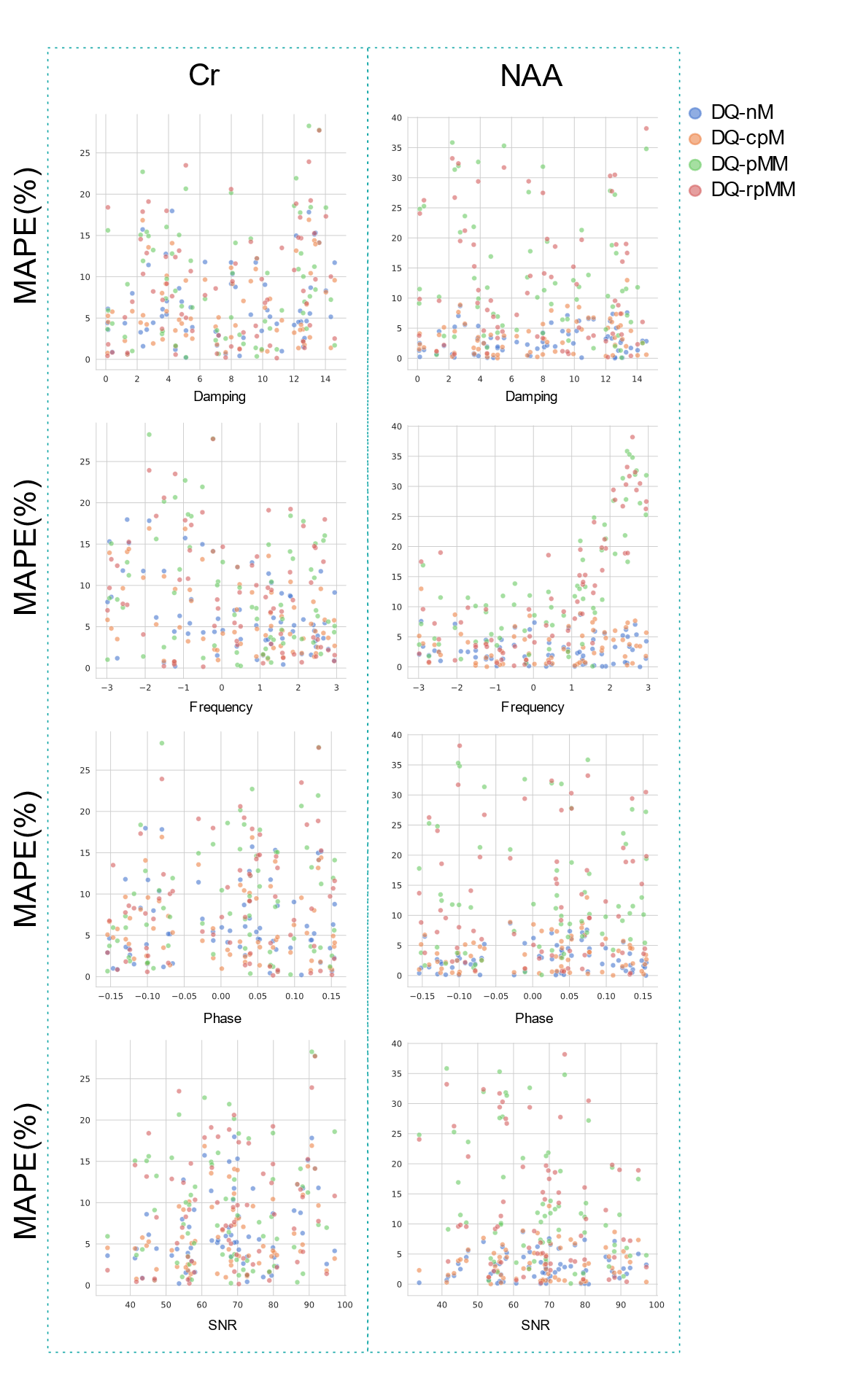

Supporting Information Figure S1. Scatter plots of the MAPEs in the estimated Cr and NAA concentration versus damping (Hz), frequency (Hz), phase (˚), and SNR of spectra.

MAPE, mean absolute percentage error; DQ-nMM, deep quantification using numeric MM; DQ-cpMM, deep quantification using numeric MM; DQ-pMM, deep quantification using parametric MM with constraint; DQ-rpMM, deep quantification using parametric MM with a regularization term.

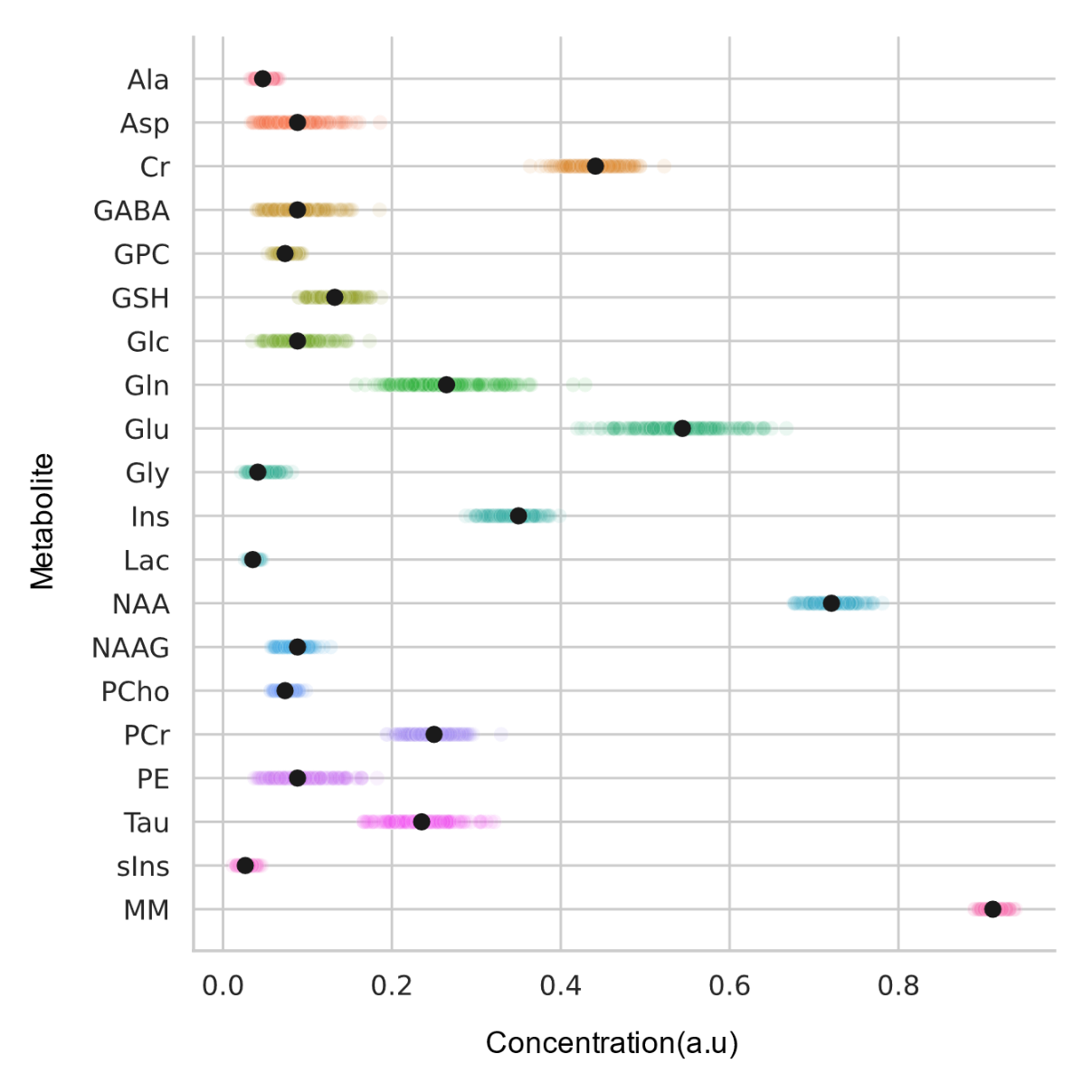

Supporting Information Figure S2. The results of MC analysis. Spectra are quantified by the DQ-nMM method. Strip plot of estimated concentration. True values are represented by black dots. DQ-nMM, deep quantification using numeric MM.

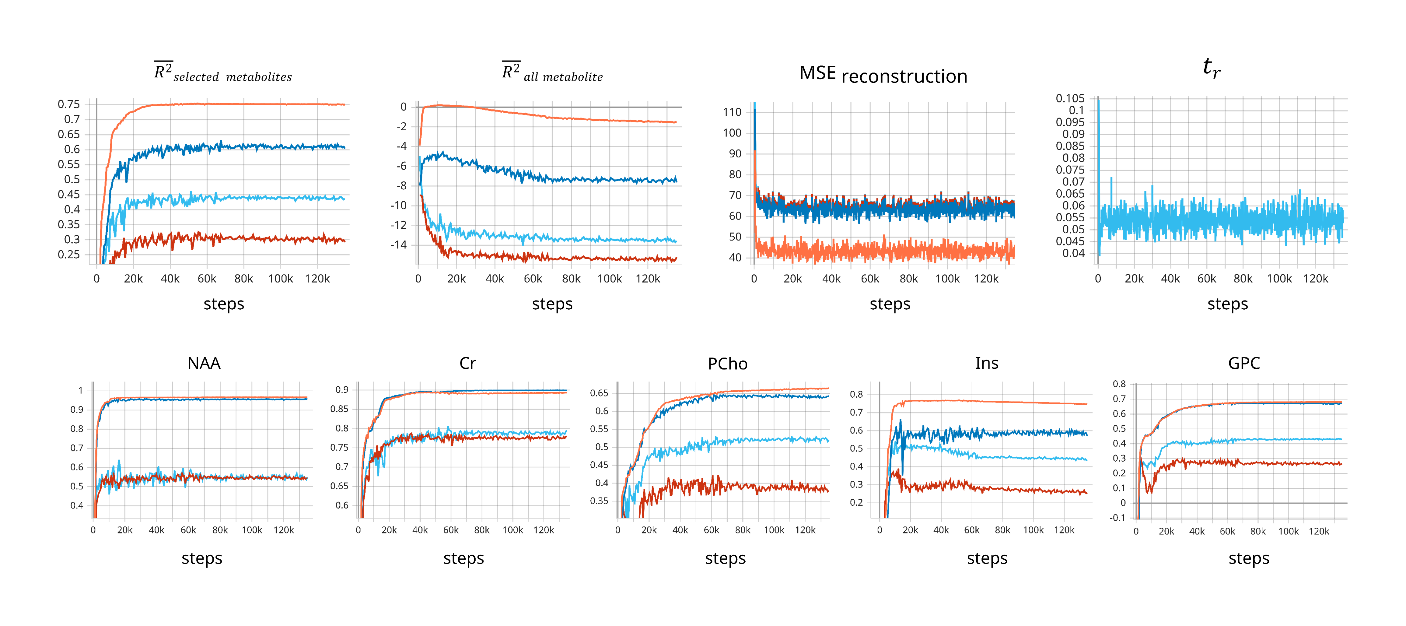

Supporting Information Figure S3. Online monitoring of metrics during training of DQ-nMM (orange), DQ-cpMM (dark blue), DQ-pMM (red), and DQ-rpMM (light blue). The mean R^2^ value calculated across the selected metabolites (Cr, GPC, Ins, NAA, PCho, Tau) and all metabolites, reconstruction loss (MSE), $t_{r}$ (the regularization term in DQ-rpMM), and the R^2^ values for NAA, Cr, PCho, Ins, and GPC against training steps are plotted.

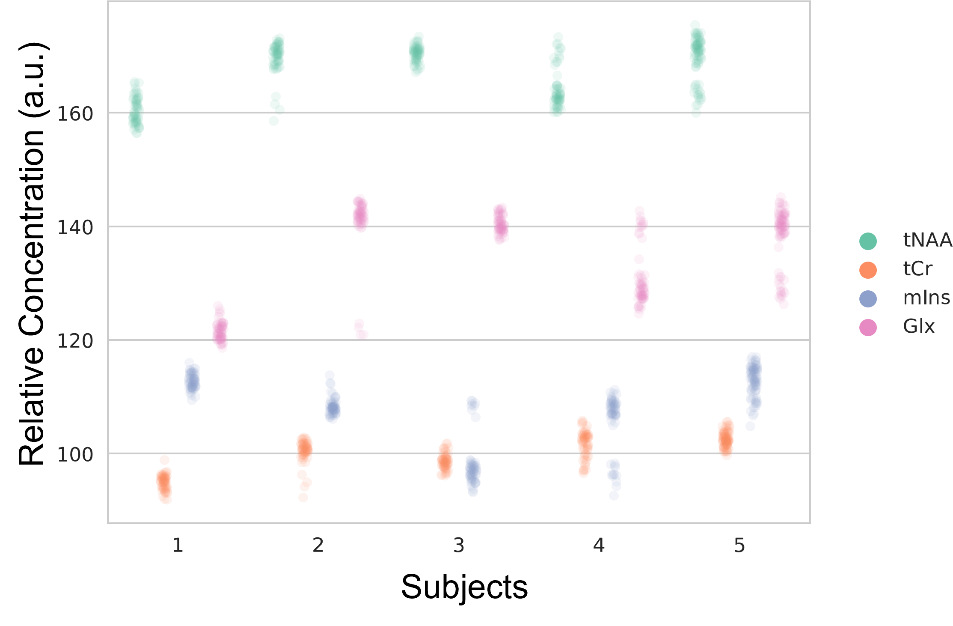

Supporting Information Figure S5. Strip plot of estimated relative concentration within subjects. The four metabolites are reported for the five subjects. Subjects are represented in the columns. Each subject has 64 augmented spectra.

Supporting Information Table S1. The details of the parametrization of the MM signal provided by the ISMRM MRS study group's fitting challenge (Marjańska et al., 2022). The MM was parametrized using AMARES (Vanhamme et al., 1997) algorithm and Lorentzian line shape for all estimated peaks. Based on our prior knowledge phase was set to 0º for all peaks.

|  | Amplitudes | Linewidths (Hz) | Frequencies (ppm) |
| --- | --- | --- | --- |
| PEAK #1 | 0.00E+00 | 5.21E+03 | -3.84E+00 |
| PEAK #2 | 3.14E-01 | 7.39E+00 | 7.64E-01 |
| PEAK #3 | 1.30E+00 | 1.06E+01 | 8.46E-01 |
| PEAK #4 | 1.62E+00 | 1.49E+01 | 9.83E-01 |
| PEAK #5 | 1.50E+00 | 1.66E+01 | 1.11E+00 |
| PEAK #6 | 6.78E-01 | 1.13E+01 | 1.53E+00 |
| PEAK #7 | 2.25E+00 | 1.29E+01 | 1.72E+00 |
| PEAK #8 | 6.74E-01 | 1.91E+01 | 1.87E+00 |
| PEAK #9 | 1.43E+00 | 2.14E+01 | 2.12E+00 |
| PEAK #10 | 0.00E+00 | 1.08E+01 | 2.41E+00 |
| PEAK #11 | 4.29E+00 | 3.17E+01 | 2.41E+00 |
| PEAK #12 | 8.10E+00 | 2.39E+01 | 2.67E+00 |
| PEAK #13 | 1.48E+00 | 2.31E+01 | 2.94E+00 |
| PEAK #14 | 1.85E+00 | 1.56E+01 | 3.24E+00 |
| PEAK #15 | 2.85E+00 | 1.91E+01 | 3.09E+00 |
| PEAK #16 | 1.52E+00 | 1.20E+01 | 3.33E+00 |
| PEAK #17 | 8.77E-01 | 1.14E+01 | 3.41E+00 |
| PEAK #18 | 1.15E+00 | 1.35E+01 | 3.50E+00 |
| PEAK #19 | 1.60E+00 | 1.20E+01 | 3.74E+00 |
| PEAK #20 | 2.21E+00 | 1.08E+01 | 3.82E+00 |

Supporting Information Table S2. Summary of the proposed network(Convolutional). Conv1d, 1D convolution over an input signal; GELU, Gaussian Error Linear Unit; Flatten, reshaping it into a one-dimensional tensor; Linear, a linear transformation to the incoming data.

| Layer (type) | Output Shape | Param # |
| --- | --- | --- |
| Conv1d-1 | [-1, 16, 510] | 176 |
| GELU-2 | [-1, 16, 510] | 0 |
| Conv1d-3 | [-1, 24, 254] | 1,176 |
| GELU-4 | [-1, 24, 254] | 0 |
| Conv1d-5 | [-1, 32, 126] | 2,336 |
| GELU-6 | [-1, 32, 126] | 0 |
| Conv1d-7 | [-1, 40, 62] | 3,880 |
| GELU-8 | [-1, 40, 62] | 0 |
| Conv1d-9 | [-1, 48, 30] | 5,808 |
| GELU-10 | [-1, 48, 30] | 0 |
| Conv1d-11 | [-1, 56, 14] | 8,120 |
| GELU-12 | [-1, 56, 14] | 0 |
| Conv1d-13 | [-1, 64, 6] | 10,816 |
| GELU-14 | [-1, 64, 6] | 0 |
| Conv1d-15 | [-1, 72, 2] | 13,896 |
| GELU-16 | [-1, 72, 2] | 0 |
| Flatten-17 | [-1, 144] | 0 |
| Linear-18 | [-1, 41] | 5,945 |
| Total params: 52,153 | | |
| Trainable params: 52,153 | | |

Supporting Information Table S3. Summary of the proposed network (MLP-Mixer). See details of each layer in (Tolstikhin et al., 2021).

| Layer (type) | | Output Shape |  | | Param # |
| --- | --- | --- | --- | --- | --- |
| Conv1d-1 | | [-1, 512, 64] |  | | 16,896 |
| Rearrange-2 | | [-1, 64, 512] |  | | 0 |
| LayerNorm-3 | | [-1, 64, 512] |  | | 1,024 |
| Rearrange-4 | | [-1, 512, 64] |  | | 0 |
| Linear-5 | | [-1, 512, 256] |  | | 16,640 |
| GELU-6 | | [-1, 512, 256] |  | | 0 |
| Dropout-7 | | [-1, 512, 256] |  | | 0 |
| Linear-8 | | [-1, 512, 64] |  | | 16,448 |
| Dropout-9 | | [-1, 512, 64] |  | | 0 |
| FeedForward-10 | | [-1, 512, 64] |  | | 0 |
| Rearrange-11 | | [-1, 64, 512] |  | | 0 |
| LayerNorm-12 | | [-1, 64, 512] |  | | 1,024 |
| Linear-13 | | [-1, 64, 2048] |  | | 1,050,624 |
| GELU-14 | | [-1, 64, 2048] |  | | 0 |
| Dropout-15 | | [-1, 64, 2048] |  | | 0 |
| Linear-16 | | [-1, 64, 512] |  | | 1,049,088 |
| Dropout-17 | | [-1, 64, 512] |  | | 0 |
| FeedForward-18 | | [-1, 64, 512] |  | | 0 |
| MixerBlock-19 | | [-1, 64, 512] |  | | 0 |
| LayerNorm-20 | | [-1, 64, 512] |  | | 1,024 |
| Rearrange-21 | | [-1, 512, 64] |  | | 0 |
| Linear-22 | | [-1, 512, 256] |  | | 16,640 |
| GELU-23 | | [-1, 512, 256] |  | | 0 |
| Dropout-24 | | [-1, 512, 256] |  | | 0 |
| Linear-25 | | [-1, 512, 64] |  | | 16,448 |
| Dropout-26 | | [-1, 512, 64] |  | | 0 |
| FeedForward-27 | | [-1, 512, 64] |  | | 0 |
| Rearrange-28 | | [-1, 64, 512] |  | | 0 |
| LayerNorm-29 | | [-1, 64, 512] |  | | 1,024 |
| Linear-30 | | [-1, 64, 2048] |  | | 1,050,624 |
| GELU-31 | | [-1, 64, 2048] |  | | 0 |
| Dropout-32 | | [-1, 64, 2048] |  | | 0 |
| Linear-33 | | [-1, 64, 512] |  | | 1,049,088 |
| Dropout-34 | | [-1, 64, 512] |  | | 0 |
| FeedForward-35 | | [-1, 64, 512] |  | | 0 |
| MixerBlock-36 | | [-1, 64, 512] |  | | 0 |
| LayerNorm-37 | | [-1, 64, 512] |  | | 1,024 |
| Rearrange-38 | | [-1, 512, 64] |  | | 0 |
| Linear-39 | | [-1, 512, 256] |  | | 16,640 |
| GELU-40 | | [-1, 512, 256] |  | | 0 |
| Dropout-41 | | [-1, 512, 256] |  | | 0 |
| Linear-42 | | [-1, 512, 64] |  | | 16,448 |
| Dropout-43 | | [-1, 512, 64] |  | | 0 |
| FeedForward-44 | | [-1, 512, 64] |  | | 0 |
| Rearrange-45 | | [-1, 64, 512] |  | | 0 |
| LayerNorm-46 | | [-1, 64, 512] |  | | 1,024 |
| Linear-47 | | [-1, 64, 2048] |  | | 1,050,624 |
| GELU-48 | | [-1, 64, 2048] |  | | 0 |
| Dropout-49 | | [-1, 64, 2048] |  | | 0 |
| Linear-50 | | [-1, 64, 512] |  | | 1,049,088 |
| Dropout-51 | | [-1, 64, 512] |  | | 0 |
| FeedForward-52 | | [-1, 64, 512] |  | | 0 |
| MixerBlock-53 | | [-1, 64, 512] |  | | 0 |
| LayerNorm-54 | | [-1, 64, 512] |  | | 1,024 |
| Rearrange-55 | | [-1, 512, 64] |  | | 0 |
| Linear-56 | | [-1, 512, 256] |  | | 16,640 |
| GELU-57 | | [-1, 512, 256] |  | | 0 |
| Dropout-58 | | [-1, 512, 256] |  | | 0 |
| Linear-59 | | [-1, 512, 64] |  | | 16,448 |
| Dropout-60 | | [-1, 512, 64] |  | | 0 |
| FeedForward-61 | | [-1, 512, 64] |  | | 0 |
| Rearrange-62 | | [-1, 64, 512] |  | | 0 |
| LayerNorm-63 | | [-1, 64, 512] |  | | 1,024 |
| Linear-64 | | [-1, 64, 2048] |  | | 1,050,624 |
| GELU-65 | | [-1, 64, 2048] |  | | 0 |
| Dropout-66 | | [-1, 64, 2048] |  | | 0 |
| Linear-67 | | [-1, 64, 512] |  | | 1,049,088 |
| Dropout-68 | | [-1, 64, 512] |  | | 0 |
| FeedForward-69 | | [-1, 64, 512] |  | | 0 |
| MixerBlock-70 | | [-1, 64, 512] |  | | 0 |
| LayerNorm-71 | | [-1, 64, 512] |  | | 1,024 |
| Rearrange-72 | | [-1, 512, 64] |  | | 0 |
| Linear-73 | | [-1, 512, 256] |  | | 16,640 |
| GELU-74 | | [-1, 512, 256] |  | | 0 |
| Dropout-75 | | [-1, 512, 256] |  | | 0 |
| Linear-76 | | [-1, 512, 64] |  | | 16,448 |
| Dropout-77 | | [-1, 512, 64] |  | | 0 |
| FeedForward-78 | | [-1, 512, 64] |  | | 0 |
| Rearrange-79 | | [-1, 64, 512] |  | | 0 |
| LayerNorm-80 | | [-1, 64, 512] |  | | 1,024 |
| Linear-81 | | [-1, 64, 2048] |  | | 1,050,624 |
| GELU-82 | | [-1, 64, 2048] |  | | 0 |
| Dropout-83 | | [-1, 64, 2048] |  | | 0 |
| Linear-84 | | [-1, 64, 512] |  | | 1,049,088 |
| Dropout-85 | | [-1, 64, 512] |  | | 0 |
| FeedForward-86 | | [-1, 64, 512] |  | | 0 |
| MixerBlock-87 | | [-1, 64, 512] |  | | 0 |
| LayerNorm-88 | | [-1, 64, 512] |  | | 1,024 |
| Rearrange-89 | | [-1, 512, 64] |  | | 0 |
| Linear-90 | | [-1, 512, 256] |  | | 16,640 |
| GELU-91 | | [-1, 512, 256] |  | | 0 |
| Dropout-92 | | [-1, 512, 256] |  | | 0 |
| Linear-93 | | [-1, 512, 64] |  | | 16,448 |
| Dropout-94 | | [-1, 512, 64] |  | | 0 |
| FeedForward-95 | | [-1, 512, 64] |  | | 0 |
| Rearrange-96 | | [-1, 64, 512] |  | | 0 |
| LayerNorm-97 | | [-1, 64, 512] |  | | 1,024 |
| Linear-98 | | [-1, 64, 2048] |  | | 1,050,624 |
| GELU-99 | | [-1, 64, 2048] |  | | 0 |
| Dropout-100 | | [-1, 64, 2048] |  | | 0 |
| Linear-101 | | [-1, 64, 512] |  | | 1,049,088 |
| Dropout-102 | | [-1, 64, 512] |  | | 0 |
| FeedForward-103 | | [-1, 64, 512] |  | | 0 |
| MixerBlock-104 | | [-1, 64, 512] |  | | 0 |
| LayerNorm-105 | | [-1, 64, 512] |  | | 1,024 |
| Rearrange-106 | | [-1, 512, 64] |  | | 0 |
| Linear-107 | | [-1, 512, 256] |  | | 16,640 |
| GELU-108 | | [-1, 512, 256] |  | | 0 |
| Dropout-109 | | [-1, 512, 256] |  | | 0 |
| Linear-110 | | [-1, 512, 64] |  | | 16,448 |
| Dropout-111 | | [-1, 512, 64] |  | | 0 |
| FeedForward-112 | | [-1, 512, 64] |  | | 0 |
| Rearrange-113 | | [-1, 64, 512] |  | | 0 |
| LayerNorm-114 | | [-1, 64, 512] |  | | 1,024 |
| Linear-115 | | [-1, 64, 2048] |  | | 1,050,624 |
| GELU-116 | | [-1, 64, 2048] |  | | 0 |
| Dropout-117 | | [-1, 64, 2048] |  | | 0 |
| Linear-118 | | [-1, 64, 512] |  | | 1,049,088 |
| Dropout-119 | | [-1, 64, 512] |  | | 0 |
| FeedForward-120 | | [-1, 64, 512] |  | | 0 |
| MixerBlock-121 | | [-1, 64, 512] |  | | 0 |
| LayerNorm-122 | | [-1, 64, 512] |  | | 1,024 |
| Rearrange-123 | | [-1, 512, 64] |  | | 0 |
| Linear-124 | | [-1, 512, 256] |  | | 16,640 |
| GELU-125 | | [-1, 512, 256] |  | | 0 |
| Dropout-126 | | [-1, 512, 256] |  | | 0 |
| Linear-127 | | [-1, 512, 64] |  | | 16,448 |
| Dropout-128 | | [-1, 512, 64] |  | | 0 |
| FeedForward-129 | | [-1, 512, 64] |  | | 0 |
| Rearrange-130 | | [-1, 64, 512] |  | | 0 |
| LayerNorm-131 | | [-1, 64, 512] |  | | 1,024 |
| Linear-132 | | [-1, 64, 2048] |  | | 1,050,624 |
| GELU-133 | | [-1, 64, 2048] |  | | 0 |
| Dropout-134 | | [-1, 64, 2048] |  | | 0 |
| Linear-135 | | [-1, 64, 512] |  | | 1,049,088 |
| Dropout-136 | | [-1, 64, 512] |  | | 0 |
| FeedForward-137 | | [-1, 64, 512] |  | | 0 |
| MixerBlock-138 | | [-1, 64, 512] |  | | 0 |
| LayerNorm-139 | | [-1, 64, 512] |  | | 1,024 |
| Linear-140 | | [-1, 41] |  | | 21,033 |
| Total params: 17,117,737 | | | | | |
| Trainable params: 17,117,737 | | | |  | |

Supporting Information Table S4. Summary of the proposed network(ConvNext). See details of each layer in (Liu et al., 2022).

| Layer (type) | Output Shape | Param # |
| --- | --- | --- |
| Conv1d-1 | [-1, 96, 256] | 864 |
| LayerNorm-2 | [-1, 96, 256] | 192 |
| Conv1d-3 | [-1, 96, 256] | 768 |
| LayerNorm-4 | [-1, 256, 96] | 192 |
| Linear-5 | [-1, 256, 384] | 37,248 |
| GELU-6 | [-1, 256, 384] | 0 |
| Linear-7 | [-1, 256, 96] | 36,960 |
| Identity-8 | [-1, 96, 256] | 0 |
| Block-9 | [-1, 96, 256] | 0 |
| Conv1d-10 | [-1, 96, 256] | 768 |
| LayerNorm-11 | [-1, 256, 96] | 192 |
| Linear-12 | [-1, 256, 384] | 37,248 |
| GELU-13 | [-1, 256, 384] | 0 |
| Linear-14 | [-1, 256, 96] | 36,960 |
| Identity-15 | [-1, 96, 256] | 0 |
| Block-16 | [-1, 96, 256] | 0 |
| Conv1d-17 | [-1, 96, 256] | 768 |
| LayerNorm-18 | [-1, 256, 96] | 192 |
| Linear-19 | [-1, 256, 384] | 37,248 |
| GELU-20 | [-1, 256, 384] | 0 |
| Linear-21 | [-1, 256, 96] | 36,960 |
| Identity-22 | [-1, 96, 256] | 0 |
| Block-23 | [-1, 96, 256] | 0 |
| LayerNorm-24 | [-1, 96, 256] | 192 |
| Conv1d-25 | [-1, 192, 64] | 73,920 |
| Conv1d-26 | [-1, 192, 64] | 1,536 |
| LayerNorm-27 | [-1, 64, 192] | 384 |
| Linear-28 | [-1, 64, 768] | 148,224 |
| GELU-29 | [-1, 64, 768] | 0 |
| Linear-30 | [-1, 64, 192] | 147,648 |
| Identity-31 | [-1, 192, 64] | 0 |
| Block-32 | [-1, 192, 64] | 0 |
| Conv1d-33 | [-1, 192, 64] | 1,536 |
| LayerNorm-34 | [-1, 64, 192] | 384 |
| Linear-35 | [-1, 64, 768] | 148,224 |
| GELU-36 | [-1, 64, 768] | 0 |
| Linear-37 | [-1, 64, 192] | 147,648 |
| Identity-38 | [-1, 192, 64] | 0 |
| Block-39 | [-1, 192, 64] | 0 |
| Conv1d-40 | [-1, 192, 64] | 1,536 |
| LayerNorm-41 | [-1, 64, 192] | 384 |
| Linear-42 | [-1, 64, 768] | 148,224 |
| GELU-43 | [-1, 64, 768] | 0 |
| Linear-44 | [-1, 64, 192] | 147,648 |
| Identity-45 | [-1, 192, 64] | 0 |
| Block-46 | [-1, 192, 64] | 0 |
| LayerNorm-47 | [-1, 192, 64] | 384 |
| Conv1d-48 | [-1, 384, 16] | 295,296 |
| Conv1d-49 | [-1, 384, 16] | 3,072 |
| LayerNorm-50 | [-1, 16, 384] | 768 |
| Linear-51 | [-1, 16, 1536] | 591,360 |
| GELU-52 | [-1, 16, 1536] | 0 |
| Linear-53 | [-1, 16, 384] | 590,208 |
| Identity-54 | [-1, 384, 16] | 0 |
| Block-55 | [-1, 384, 16] | 0 |
| Conv1d-56 | [-1, 384, 16] | 3,072 |
| LayerNorm-57 | [-1, 16, 384] | 768 |
| Linear-58 | [-1, 16, 1536] | 591,360 |
| GELU-59 | [-1, 16, 1536] | 0 |
| Linear-60 | [-1, 16, 384] | 590,208 |
| Identity-61 | [-1, 384, 16] | 0 |
| Block-62 | [-1, 384, 16] | 0 |
| Conv1d-63 | [-1, 384, 16] | 3,072 |
| LayerNorm-64 | [-1, 16, 384] | 768 |
| Linear-65 | [-1, 16, 1536] | 591,360 |
| GELU-66 | [-1, 16, 1536] | 0 |
| Linear-67 | [-1, 16, 384] | 590,208 |
| Identity-68 | [-1, 384, 16] | 0 |
| Block-69 | [-1, 384, 16] | 0 |
| Conv1d-70 | [-1, 384, 16] | 3,072 |
| LayerNorm-71 | [-1, 16, 384] | 768 |
| Linear-72 | [-1, 16, 1536] | 591,360 |
| GELU-73 | [-1, 16, 1536] | 0 |
| Linear-74 | [-1, 16, 384] | 590,208 |
| Identity-75 | [-1, 384, 16] | 0 |
| Block-76 | [-1, 384, 16] | 0 |
| Conv1d-77 | [-1, 384, 16] | 3,072 |
| LayerNorm-78 | [-1, 16, 384] | 768 |
| Linear-79 | [-1, 16, 1536] | 591,360 |
| GELU-80 | [-1, 16, 1536] | 0 |
| Linear-81 | [-1, 16, 384] | 590,208 |
| Identity-82 | [-1, 384, 16] | 0 |
| Block-83 | [-1, 384, 16] | 0 |
| Conv1d-84 | [-1, 384, 16] | 3,072 |
| LayerNorm-85 | [-1, 16, 384] | 768 |
| Linear-86 | [-1, 16, 1536] | 591,360 |
| GELU-87 | [-1, 16, 1536] | 0 |
| Linear-88 | [-1, 16, 384] | 590,208 |
| Identity-89 | [-1, 384, 16] | 0 |
| Block-90 | [-1, 384, 16] | 0 |
| Conv1d-91 | [-1, 384, 16] | 3,072 |
| LayerNorm-92 | [-1, 16, 384] | 768 |
| Linear-93 | [-1, 16, 1536] | 591,360 |
| GELU-94 | [-1, 16, 1536] | 0 |
| Linear-95 | [-1, 16, 384] | 590,208 |
| Identity-96 | [-1, 384, 16] | 0 |
| Block-97 | [-1, 384, 16] | 0 |
| Conv1d-98 | [-1, 384, 16] | 3,072 |
| LayerNorm-99 | [-1, 16, 384] | 768 |
| Linear-100 | [-1, 16, 1536] | 591,360 |
| GELU-101 | [-1, 16, 1536] | 0 |
| Linear-102 | [-1, 16, 384] | 590,208 |
| Identity-103 | [-1, 384, 16] | 0 |
| Block-104 | [-1, 384, 16] | 0 |
| Conv1d-105 | [-1, 384, 16] | 3,072 |
| LayerNorm-106 | [-1, 16, 384] | 768 |
| Linear-107 | [-1, 16, 1536] | 591,360 |
| GELU-108 | [-1, 16, 1536] | 0 |
| Linear-109 | [-1, 16, 384] | 590,208 |
| Identity-110 | [-1, 384, 16] | 0 |
| Block-111 | [-1, 384, 16] | 0 |
| LayerNorm-112 | [-1, 384, 16] | 768 |
| Conv1d-113 | [-1, 768, 4] | 1,180,416 |
| Conv1d-114 | [-1, 768, 4] | 6,144 |
| LayerNorm-115 | [-1, 4, 768] | 1,536 |
| Linear-116 | [-1, 4, 3072] | 2,362,368 |
| GELU-117 | [-1, 4, 3072] | 0 |
| Linear-118 | [-1, 4, 768] | 2,360,064 |
| Identity-119 | [-1, 768, 4] | 0 |
| Block-120 | [-1, 768, 4] | 0 |
| Conv1d-121 | [-1, 768, 4] | 6,144 |
| LayerNorm-122 | [-1, 4, 768] | 1,536 |
| Linear-123 | [-1, 4, 3072] | 2,362,368 |
| GELU-124 | [-1, 4, 3072] | 0 |
| Linear-125 | [-1, 4, 768] | 2,360,064 |
| Identity-126 | [-1, 768, 4] | 0 |
| Block-127 | [-1, 768, 4] | 0 |
| Conv1d-128 | [-1, 768, 4] | 6,144 |
| LayerNorm-129 | [-1, 4, 768] | 1,536 |
| Linear-130 | [-1, 4, 3072] | 2,362,368 |
| GELU-131 | [-1, 4, 3072] | 0 |
| Linear-132 | [-1, 4, 768] | 2,360,064 |
| Identity-133 | [-1, 768, 4] | 0 |
| Block-134 | [-1, 768, 4] | 0 |
| Linear-135 | [-1, 41] | 31,529 |
| Total params: 27,561,449 | | |
| Trainable params: 27,561,449 |  |  |
| Non-trainable params: 0 | | |

Supporting Information Table S5. Summary of the proposed network(Fully connected). GELU, Gaussian Error Linear Unit; Flatten, reshaping it into a one-dimensional tensor; Linear, a linear transformation to the incoming data.

| Layer (type) | Output Shape | Param # |
| --- | --- | --- |
| Linear-1 | [-1, 2, 768] | 787,200 |
| GELU-2 | [-1, 2, 768] | 0 |
| Linear-3 | [-1, 2, 512] | 393,728 |
| GELU-4 | [-1, 2, 512] | 0 |
| Linear-5 | [-1, 2, 256] | 131,328 |
| GELU-6 | [-1, 2, 256] | 0 |
| Linear-7 | [-1, 2, 128] | 32,896 |
| GELU-8 | [-1, 2, 128] | 0 |
| Linear-9 | [-1, 2, 64] | 8,256 |
| GELU-10 | [-1, 2, 64] | 0 |
| Flatten-11 | [-1, 128] | 0 |
| Linear-12 | [-1, 41] | 5,289 |
| Total params: 1,358,697 | |  |
| Trainable params: 1,358,697 | |  |

Supporting Information Text S1: The detailed model of macromolecules

The macromolecules term ($A_{b}B\left( t \right)$) in Eq. 1 can be expressed as

$$(\sum_{n=1}^{N} A_{n}B_{n}\left( t \right) e^{\left( 2i\pi f_{n} \right)t})e^{\alpha t}$$

where $B_{n}\left( t \right)$, $A_{bn}$ and $f_{n},$ are the signal, the scaling factor, and the frequency shift of the n-th MM component, respectively, and $\alpha$ is the global damping factor.

Supporting Information Text S2: Bayesian hyper-parameterization

The Bayesian Optimization HyperBand method(Falkner et al., 2018) from the Tune framework(Liaw et al., 2018) was utilized for the optimization of the encoder's hyperparameters. A total of 30 model trials were conducted, with the validation loss being monitored (MSE). The training was conducted for 50 epochs, and Tune's built-in early termination criteria were used(Liaw et al., 2018).

Based on prior funding, we narrowed our parameters to only the four most significant parameters, which are as follows:

The learning rate: 5e-5 (fixed)

The depth of the encoder, i.e., number of convolutional blocks: {4, 6, 8}

$\lambda$: (2, 7)

Batch size: {32, 128}

Kernel size of convolotion layer: {5, 7}

Finally, the Bayesian Optimization HyperBand method explored the parameter space and identified the model with the best tuning and the least validation loss. The following table summarizes the tuned parameters:

The learning rate: 5e-05

The depth of the encoder: 4

Batch size:32

$\lambda$: 3.94

Kernel size of convolotion layer: 7

It should be noted that since the network was trained without supervision, the smallest MSE loss does not necessarily correspond to the smallest prediction error. We observed the R_2_ (coefficient of determination) value of predicted frequency and phase shifts in each mini-batch during training(Supplemental Fig. 4.) on the simulated dataset with known shifts. We further optimized the encoder's settings to get the highest R_2_. Fine-tuned parameters are reported in the implementation details section.

Falkner, S., Klein, A., Hutter, F., 2018. BOHB: Robust and Efficient Hyperparameter Optimization at Scale.

Liaw, R., Liang, E., Nishihara, R., Moritz, P., Gonzalez, J.E., Stoica, I., 2018. Tune: A Research Platform for Distributed Model Selection and Training.

Liu, Z., Mao, H., Wu, C.-Y., Feichtenhofer, C., Darrell, T., Xie, S., 2022. A ConvNet for the 2020s. arXiv. https://doi.org/10.48550/arxiv.2201.03545

Marjańska, M., Deelchand, D.K., Kreis, R., Alger, J.R., Bolan, P.J., Borbath, T., Boumezbeur, F., Fernandes, C.C., Coello, E., Nagraja, B.H., Považan, M., Ratiney, H., Sima, D., Starčuková, J., Soher, B.J., Wilson, M., van Asten, J.J.A., 2022. Results and interpretation of a fitting challenge for MR spectroscopy set up by the MRS study group of ISMRM. Magn. Reson. Med. 87, 11–32. https://doi.org/10.1002/mrm.28942

Tolstikhin, I., Houlsby, N., Kolesnikov, A., Beyer, L., Zhai, X., Unterthiner, T., Yung, J., Steiner, A., Keysers, D., Uszkoreit, J., Lucic, M., Dosovitskiy, A., 2021. MLP-Mixer: An all-MLP Architecture for Vision. arXiv. https://doi.org/10.48550/arXiv.2105.01601

Vanhamme, L., Van Den Boogaart, A., Van Huffel, S., 1997. Improved Method for Accurate and Efficient Quantification of MRS Data with Use of Prior Knowledge. J. Magn. Reson. 129, 35–43. https://doi.org/10.1006/jmre.1997.1244
